# A neurotransmitter atlas of *C. elegans* males and hermaphrodites

**DOI:** 10.1101/2023.12.24.573258

**Authors:** Chen Wang, Berta Vidal, Surojit Sural, Curtis Loer, G. Robert Aguilar, Daniel M. Merritt, Itai Antoine Toker, Merly C. Vogt, Cyril Cros, Oliver Hobert

## Abstract

Mapping neurotransmitter identities to neurons is key to understanding information flow in a nervous system. It also provides valuable entry points for studying the development and plasticity of neuronal identity features. In the *C. elegans* nervous system, neurotransmitter identities have been largely assigned by expression pattern analysis of neurotransmitter pathway genes that encode neurotransmitter biosynthetic enzymes or transporters. However, many of these assignments have relied on multicopy reporter transgenes that may lack relevant *cis*-regulatory information and therefore may not provide an accurate picture of neurotransmitter usage. We analyzed the expression patterns of 16 CRISPR/Cas9-engineered knock-in reporter strains for all main types of neurotransmitters in *C. elegans* (glutamate, acetylcholine, GABA, serotonin, dopamine, tyramine, and octopamine) in both the hermaphrodite and the male. Our analysis reveals novel sites of expression of these neurotransmitter systems within both neurons and glia, as well as non-neural cells. The resulting expression atlas defines neurons that may be exclusively neuropeptidergic, substantially expands the repertoire of neurons capable of co-transmitting multiple neurotransmitters, and identifies novel neurons that uptake monoaminergic neurotransmitters. Furthermore, we also observed unusual co-expression patterns of monoaminergic synthesis pathway genes, suggesting the existence of novel monoaminergic transmitters. Our analysis results in what constitutes the most extensive whole-animal-wide map of neurotransmitter usage to date, paving the way for a better understanding of neuronal communication and neuronal identity specification in *C. elegans*.

## INTRODUCTION

Understanding information processing in the brain necessitates the generation of precise maps of neurotransmitter deployment. Moreover, comprehending synaptic wiring diagrams is contingent upon decoding the nature of signaling events between anatomically connected neurons. Mapping of neurotransmitter identities onto individual neuron classes also presents a valuable entry point for studying how neuronal identity features become genetically specified during development and potentially modified in response to specific external factors (such as the environment) or internal factors (such as sexual identity or neuronal activity patterns).

The existence of complete synaptic wiring diagrams of the compact nervous system of male and hermaphrodite *C. elegans* nematodes raises questions about the molecular mechanisms by which individual neurons communicate with each other. *C. elegans* employs the main neurotransmitter systems that are used throughout the animal kingdom, including acetylcholine, glutamate, γ-aminobutyric acid (GABA), and several monoamines (Sulston *et al*. 1975; Horvitz *et al*. 1982; Loer and Kenyon 1993; Mcintire *et al*. 1993; Duerr *et al*. 1999; Lee *et al*. 1999; Duerr *et al*. 2001; Alkema *et al*. 2005; Duerr *et al*. 2008; Serrano-Saiz *et al*. 2013; Pereira *et al*. 2015; Gendrel *et al*. 2016; Serrano-Saiz *et al*. 2017b)(**Fig. 1A**). Efforts to map these neurotransmitter systems to individual cell types throughout the entire nervous system have a long history, beginning with the use of chemical stains that directly detected a given neurotransmitter (dopamine)(Sulston *et al*. 1975), followed by antibody staining of neurotransmitter themselves (serotonin and GABA)(Horvitz *et al*. 1982; Mcintire *et al*. 1993) or antibody stains of biosynthetic enzymes or neurotransmitter vesicular transporters (acetylcholine and monoamines)(LOER AND KENYON 1993; Duerr *et al*. 1999; Duerr *et al*. 2001; Alkema *et al*. 2005; Duerr *et al*. 2008)(see **Fig. 1A** for an overview of these enzymes and transporters).

**Figure 1.**
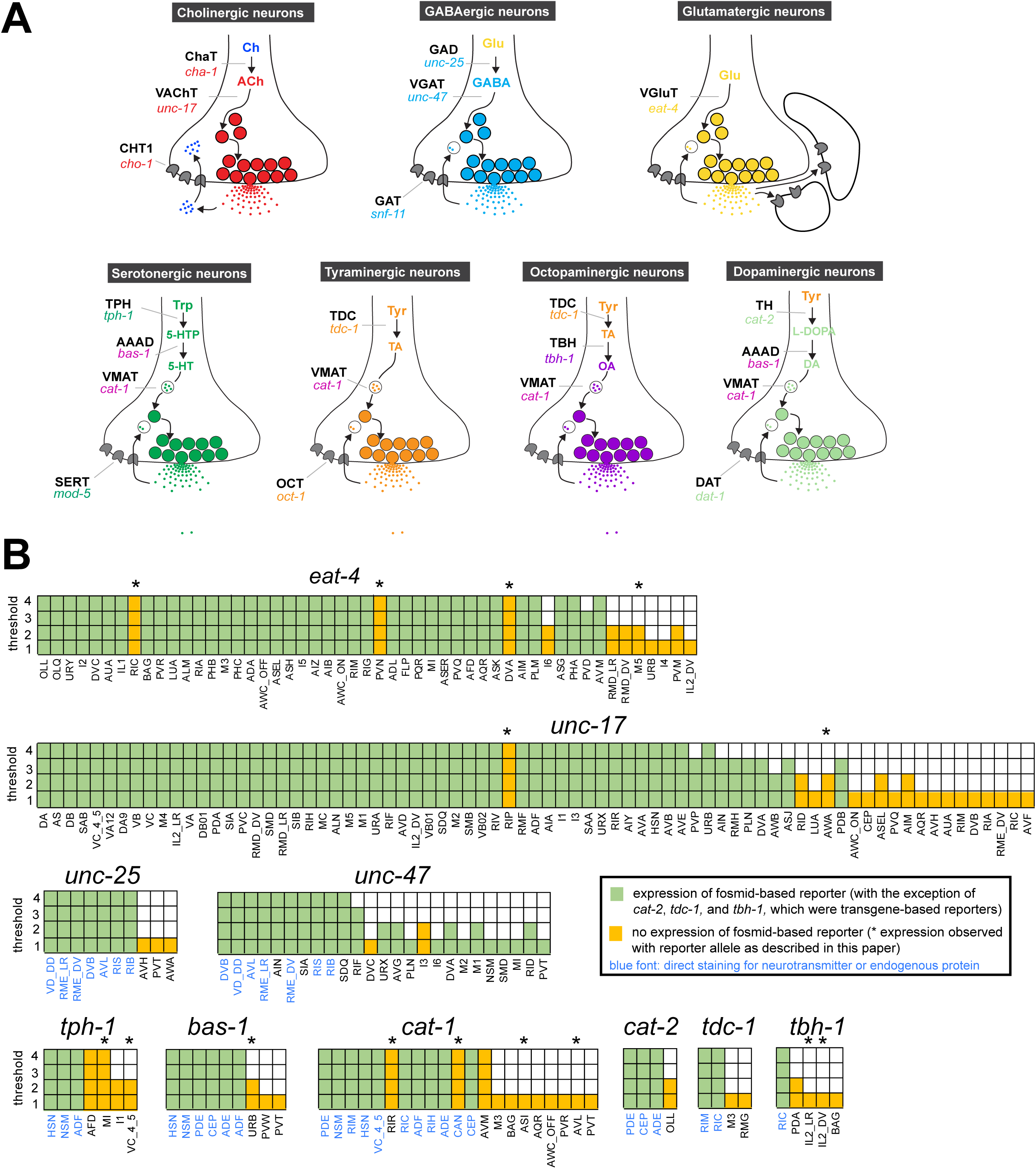
Background on genes examined in this paper. **(A)** Neurotransmitter synthesis and transport pathways. TH = tyrosine hydroxylase; TDC = tyrosine decarboxylase; TBH = tyramine beta hydroxylase; TPH = tryptophan hydroxylase; GAD = glutamic acid decarboxylase; AAAD = aromatic amino acid decarboxylase; VMAT = vesicular monoamine transporter; VAChT = vesicular acetylcholine transporter; VGAT = vesicular GABA transporter. Taken and modified from (HOBERT 2013). **(B)** Graphic comparison of scRNA expression data and previously reported reporter expression data. See **Table S1** for a more comprehensive version that includes expression of reporter genes in cells that show no scRNA transcripts. Note that scRNA expression values for *eat-4* and *unc-47* can be unreliable because they were overexpressed to isolate individual neurons for scRNA analysis (Taylor *et al*. 2021).

While these early approaches proved successful in revealing neurotransmitter identities, they displayed several technical limitations. Since neurotransmitter-synthesizing or - transporting proteins primarily localize to neurites, the cellular identity of expressing cells (usually determined by assessing cell body position) often could not be unambiguously established in several, particularly cell- and neurite-dense regions of the nervous system. One example concerns cholinergic neurons, which are defined by the expression of the vesicular acetylcholine transporter UNC-17/VAChT and choline acetyltransferase CHA-1/ChAT. While mainly neurite-localized UNC-17 and CHA-1 antibody staining experiments could identify a subset of cholinergic neurons (Duerr *et al*. 2001; Duerr *et al*. 2008), many remained unidentified (Pereira *et al*. 2015). In addition, for GABA-producing neurons, it became apparent that antibody-based GABA detection was dependent on staining protocols, leading to the identification of “novel” anti-GABA-positive neurons, i.e. GABAergic neurons, more than 20 years after the initial description of GABAergic neurons (Mcintire *et al*. 1993; Gendrel *et al*. 2016).

An alternative approach to mapping neurotransmitter usage has been the use of reporter transgenes. This approach has the significant advantage of allowing the fluorophore to either fill the entire cytoplasm of a cell or to be targeted to the nucleus, thereby facilitating neuron identification. However, one shortcoming of transgene-based reporter approaches is that one cannot be certain that a chosen genomic region, fused to a reporter gene, indeed contains all *cis*-regulatory elements of the respective locus. In fact, the first report that described the expression of the vesicular glutamate transporter EAT-4, the key marker for glutamatergic neuron identity, largely underestimated the number of *eat-4/VLGUT-*positive and, hence, glutamatergic neurons (Lee *et al*. 1999). The introduction of fosmid-based reporter transgenes has largely addressed such concerns, as these reporters, with their 30-50 Kb size, usually cover entire intergenic regions (SAROV *et al*. 2012). Indeed, such fosmid-based reporters have been instrumental in describing the entire *C. elegans* glutamatergic nervous system, defined by the expression of *eat-4/VGLUT* (Serrano-Saiz *et al*. 2013), as well as supposedly the complete set of cholinergic (Pereira *et al*. 2015) and GABAergic neurons (Gendrel *et al*. 2016).

However, even fosmid-based reporters may not be the final word. In theory, they may still miss distal *cis*-regulatory elements. Moreover, the multicopy nature of transgenes harbors the risk of overexpression artifacts, such as the titrating of rate-limiting negative regulatory mechanisms. Also, RNAi-based silencing mechanisms triggered by the multicopy nature of transgenic reporter arrays have the potential to dampen the expression of reporter arrays (NANCE AND FROKJAER-JENSEN 2019). One way to get around these limitations, while still preserving the advantages of reporter gene approaches, is to generate reporter alleles in which an endogenous locus is tagged with a reporter cassette, using CRISPR/Cas9 genome engineering. Side-by-side comparisons of fosmid-based reporter expression patterns with those of knock-in reporter alleles indeed revealed several instances of discrepancies in expression patterns of homeobox genes (Reilly *et al*. 2022).

An indication that previous neurotransmitter assignments may not have been complete was provided by recent single-cell RNA (scRNA) transcriptomic analyses of the hermaphrodite nervous system by the CeNGEN consortium (Taylor *et al*. 2021). As we describe in this paper in more detail, transcripts for several neurotransmitter-synthesizing enzymes or transporters were detected in a few cells beyond those previously described to express the respective genes. This motivated us to use CRISPR/Cas9 engineering to fluorescently tag a comprehensive panel of genetic loci that code for neurotransmitter-synthesizing, -transporting, and -uptaking proteins (“neurotransmitter pathway genes”). Using the landmark strain NeuroPAL for neuron identification (Yemini *et al*. 2021), we identified novel sites of expression of most neurotransmitter pathway genes. Furthermore, we used these reagents to expand and refine neurotransmitter maps of the entire nervous system of the *C. elegans* male, which contains almost 30% more neurons than the nervous system of the hermaphrodite yet lacks a reported scRNA transcriptome atlas. Together with the NeuroPAL cell-identification tool, these reporter alleles allowed us to substantially improve the previously described neurotransmitter map of the male nervous system (Serrano-Saiz *et al*. 2017b). Our analysis provides insights into the breadth of usage of each individual neurotransmitter system, reveals instances of co-transmitter use, indicates the existence of neurons that may entirely rely on neuropeptides instead of classic neurotransmitters, reveals sexual dimorphisms in neurotransmitter usage, and suggests the likely existence of presently unknown neurotransmitters.

## MATERIALS AND METHODS

### Transgenic reporter strains

Knock-in reporter alleles were generated either by SunyBiotech (*syb* alleles) or in-house (*ot* alleles) using CRISPR/Cas9 genome engineering. Most genes were tagged with a nuclear-targeted *gfp* sequence (*gfp* fused to *his-44*, a histone *h2b* gene) at the 3’ end of the locus to capture all isoforms, except *tdc-1* which was tagged at the 5’ end. For *unc-25*, both isoforms were individually tagged since a single tag would not capture both. Transgene schematics are shown in **Fig. 2**.

**Figure 2.**
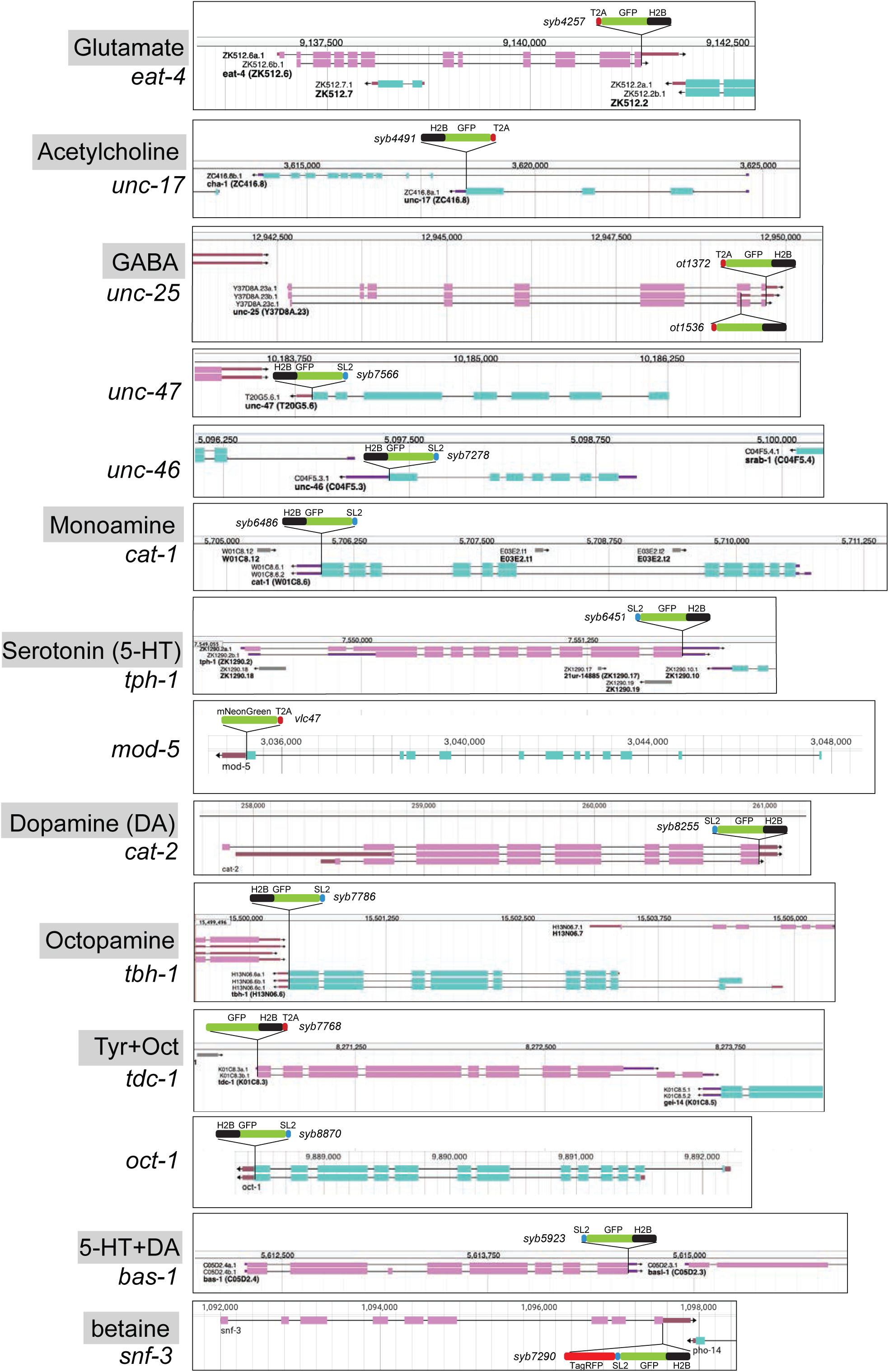
Schematics of reporter knock-in alleles. Reporter alleles were generated by CRISPR/Cas9 genome engineering. The SL2 or T2A-based separation of the reporter from the coding sequence of the respective loci enables targeting of the reporter to the nucleus (via the H2B tag), which in turn facilitates identification of the cell expressing a given reporter. Genome schematics are from WormBase (Davis *et al*. 2022).

Reporter alleles generated in this study:

*unc-25(ot1372[unc-25a.1c.1::t2a:gfp::h2b]) III*

*unc-25(ot1536[unc-25b.1::t2a::gfp::h2b]) III*

*unc-46(syb7278[unc-46::sl2::gfp:h2b]) V*

*unc-47(syb7566[unc-47::sl2::gfp::h2b]) III*

*cat-1(syb6486[cat-1::sl2::gfp::h2b]) X*

*tph-1(syb6451[tph-1::sl2::gfp::h2b]) II*

*tbh-1(syb7786[tbh-1::sl2::gfp::h2b]) X*

*tdc-1(syb7768[gfp::linker::h2b::t2a::tdc-1]) II*

*cat-2(syb8255[cat-2::sl2::gfp::h2b]) II*

*snf-3(syb7290[snf-3::TagRFP::sl2::gfp::h2b]) II*

*oct-1(syb8870[oct-1::sl2::gfp::h2b]) I*

*hdl-1(syb1048[hdl-1::gfp]) IV*

*hdl-1(syb4208[hdl-1::t2a::3xnls::cre]) IV*

Since we did not detect fluorophore signals in the *hdl-1(syb1048[hdl-1::gfp])* strain, we attempted to amplify low level signals, by inserting Cre recombinase at the C-terminus of the *hdl-1* locus (*hdl-1(syb4208[hdl-1::t2a::3xnls::cre])*). We crossed this strain to the recently published “Flexon” strain (*arTi361[rps-27p::gfp"flexon"-h2b::unc-54-3’UTR]*) (SHAFFER AND GREENWALD 2022). Even low expression of *hdl-1* should have led to Cre-mediated excision of the flexon stop cassette, which is designed to abrogate gene expression by a translational stop and frameshift mutation, and subsequently can result in strong and sustained *gfp* expression under the control of the *rps-27* promoter and thereby providing information about cell-specific *hdl-1* expression. However, no robust, consistent reporter expression was seen in *hdl-1(syb4208[hdl-1::t2a::3xnls::cre]); arTi361[rps-27p::gfp"flexon"-h2b::unc-54-3’UTR]* animals.

Three of the reporter alleles that we generated were already previously examined in specific cellular contexts:

*unc-17(syb4491[unc-17::t2a::gfp:h2b]) IV* (Vidal *et al*. 2022)

*eat-4(syb4257[eat-4::t2a::gfp::h2b]) III* (Vidal *et al*. 2022)

*bas-1(syb5923[bas-1::sl2::gfp::h2b]) III* (Yu *et al*. 2023)

One of the reporter alleles was obtained from the Caenorhabditis Genetics Center (CGC):

*mod-5(vlc47[mod-5::t2a::mNeonGreen]) I* (MAICAS et al. 2021)

### Microscopy and image processing

For adult animal imaging, 15-25 (exact number depending on the difficulty of neuron ID) same-sex L4 worms were grouped on NGM plates 6-9 hours prior to imaging to control for accurate staging and avoid mating. Young adult worms were then anesthetized using 50—100 mM sodium azide and mounted on 5% agarose pads on glass slides. Z-stack images were acquired with ZEN software using Zeiss confocal microscopes LSM880 and LSM980 or a Zeiss Axio Imager Z2 and processed with ZEN software or FIJI (Schindelin *et al*. 2012) to create orthogonal projections. Brightness and contrast, and in some cases gamma values, were adjusted to illustrate dim expression and facilitate neuron identification.

### Neuron class and cell type identification

Neuron classes were identified by crossing the *gfp* reporter alleles with the landmark strain “NeuroPAL” (allele *otIs669* or *otIs696*, for bright reporters and dim reporters, respectively) and following published protocols (Tekieli *et al*. 2021; Yemini *et al*. 2021) (also see “lab resources” at hobertlab.org). For neuron identification of the *eat-4(syb4257)*, *unc-46(syb7278)*, and *unc-47(syb7566)* reporter alleles in hermaphrodites, the reporter alleles were also crossed into the fosmid-based reporter transgenes of the same gene [*eat-4(otIs518)*, *unc-46(otIs568)*, *and unc-47(otIs564)*] as a “first-pass” to identify potential non-overlapping expression of the two alleles. For *tph-1(syb6451)* analysis, an *eat-4* fosmid-based reporter (*otIs518*) was also used. For identification of VC4, VC5, HSN, and uv1, an *ida-1p::mCherry* integrant (LX2478, *lin-15(n765ts) X; vsls269[ida-1::mCherry]*) was also used in some cases (Fernandez *et al*. 2020). For phasmid neurons, dye-filling with DiD (Thermo Fisher Scientific) was sometimes used to confirm neuron ID. For glial expression, a panglial reporter *otIs870[mir-228p::3xnls::TagRFP]* was used. For hypodermal cells identification, a *dpy-7p::mCherry* reporter *stIs10166 [dpy-7p::his-24::mCherry + unc-119(+)]* was used. (Liu *et al*. 2009)

## RESULTS

### Comparing CeNGEN scRNA data to reporter gene data

To investigate the neurotransmitter identity of neurons throughout the entire *C. elegans* nervous system of both sexes, we consider here the expression pattern of the following 15 genetic loci (see also **Fig. 1A**):

a. *eat-4/VGLUT*: expression of the vesicular glutamate transporter is alone sufficient to define glutamatergic neuron identity (Lee *et al*. 1999; Serrano-Saiz *et al*. 2013).
b. *unc-17/VAChT*: expression of the vesicular acetylcholine transporter, located in an operon together with the acetylcholine-synthesizing gene *cha-1/ChAT* (Alfonso *et al*. 1994), defines cholinergic neurons (Duerr *et al*. 2001; Duerr *et al*. 2008; Pereira *et al*. 2015).
c. *unc-25/GAD, unc-47/VGAT* and its sorting co-factor *unc-46*/*LAMP*: expression of these three genes defines neurons that synthesize and release GABA (Mcintire *et al*. 1993; Mcintire *et al*. 1997; Jin *et al*. 1999; Schuske *et al*. 2007; Gendrel *et al*. 2016). Additional neurons that we classify as GABAergic are those that do not synthesize GABA (*unc-25/GAD*-negative), but take up GABA from other neurons (based on anti-GABA antibody staining) and are expected to release GABA based on *unc-47/VGAT* expression (Gendrel *et al*. 2016)*. unc-47/VGAT* expression without any evidence of GABA synthesis or uptake (*unc-25/GAD-* and anti-GABA-negative) is indicative of an unknown transmitter being present in these cells and utilizing *unc-47/VGAT* for vesicular secretion.
d. *tph-1/TPH* and *bas-1/AAAD*: the co-expression of these two biosynthetic enzymes, together with the co-expression of the monoamine vesicular transporter *cat-1/VMAT,* defines all serotonin-synthesizing and -releasing neurons (**Fig. 1A**)(Horvitz *et al*. 1982; Duerr *et al*. 1999; Sze *et al*. 2000; HARE AND LOER 2004).
e. *cat-2/TH* and *bas-1/AAAD*: the co-expression of these two biosynthetic enzymes, together with the co-expression of the monoamine vesicular transporter *cat-1/VMAT,* defines all dopamine-synthesizing and -releasing neurons (**Fig. 1A**)(Sulston *et al*. 1975; Duerr *et al*. 1999; LINTS AND EMMONS 1999; HARE AND LOER 2004).
f. *tdc-1/TDC*: defines, together with *cat-1/VMAT,* all tyramine-synthesizing and -releasing neurons (**Fig. 1A**)(Alkema *et al*. 2005).
g. *tbh-1/TBH*: expression of this gene, in combination with that of *tdc-1/TDC* and *cat-1/VMAT*, defines octopamine-synthesizing and -releasing neurons (**Fig. 1A**) (Alkema *et al*. 2005).
h. *cat-1/VMAT*: expression of this vesicular monoamine transporter defines all four above-mentioned monoaminergic neurons (serotonin, dopamine, tyramine, octopamine)(Duerr *et al*. 1999), but as described and discussed below, it may also define additional sets of monoaminergic neurons.
i. *hdl-1/AAAD*: *hdl-1,* a previously uncharacterized gene, encodes the only other AAAD with sequence similarity to the *bas-1* and *tdc-1* AAAD enzymes that produce other *bona fide* monoamines (**Fig. S1**)(HARE AND LOER 2004). *hdl-1* expression may therefore, in combination with *cat-1/VMAT,* identify neurons that produce and release trace amines of unknown identity.
j. *snf-3/BGT1/SLC6A12*: this gene encodes the functionally validated orthologue of the vertebrate betaine uptake transporter SLC6A12 (i.e. BGT1)(Peden *et al*. 2013). In combination with the expression of *cat-1/VMAT,* which synaptically transports betaine (Hardege *et al*. 2022)*, snf-3* expression may identify neurons that utilize betaine as a synaptically released neurotransmitter to gate betaine-gated ion channels, such as ACR-23 (Peden *et al*. 2013) or LGC-41 (Hardege *et al*. 2022).
k. *mod-5/SERT*: this gene codes for the functionally validated orthologue of the vertebrate serotonin uptake transporter SERT (Ranganathan *et al*. 2001), which defines neurons that take up 5-HT independently of their ability to synthesize 5-HT and, depending on their expression of *cat-1/VMAT*, may either re-utilize serotonin for synaptic signaling or serve as 5-HT clearance neurons.
l. *oct-1/OCT*: this gene encodes the closest representative of the OCT subclass of SLC22 organic anion transporters (Zhu *et al*. 2015), several members of which are selective uptake transporters of tyramine (Breidert *et al*. 1998; Berry *et al*. 2016). Its expression or function in the nervous system had not previously been analyzed in *C. elegans*.

For all these 15 genetic loci, we compared scRNA transcriptome data from the CeNGEN scRNA atlas (at all 4 available stringency levels (Taylor *et al*. 2021)) to previously published reporter and antibody staining data. As shown in **Fig. 1B** and **Table S1**, such comparisons reveal the following: (1) scRNA data support the expression of genes in the vast majority of neurons in which those genes were found to be expressed with previous reporter gene approaches. In most cases, this is true even at the highest threshold levels for scRNA detection. (2) Vice versa, reporter gene expression supports scRNA transcriptome data for a specific neurotransmitter system in the great majority of cells. (3) In spite of this congruence, there were several discrepancies between reporter data and scRNA data. Generally, while valuable, scRNA transcriptome data cannot be considered the final word for any gene expression pattern assignments. Lack of detection of transcripts could be a sensitivity issue and, conversely, the presence of transcripts does not necessarily indicate that the respective protein is generated, due to the possibility of posttranscriptional regulation.

Hence, to consolidate and further improve neurotransmitter identity assignment throughout the entire *C. elegans* nervous system, and to circumvent potential limitations of multicopy, fosmid-based reporter transgenes on which previous neurotransmitter assignments have been based, we engineered and examined expression patterns of 16 knock-in reporter alleles of neurotransmitter synthesis, vesicular transport, and uptake loci listed above (**Fig. 1**, **Fig. 2**). For *unc-17* and *eat-4,* we knocked-in a *t2a::gfp::h2b* (*his-44*) cassette right before the stop codon of the respective gene. For *unc-25*, we created two knock-in alleles with the *t2a::gfp::h2b* (*his-44*) cassette tagging isoforms a.1/c.1 and b.1 separately. For *tdc-1,* a *gfp::h2b::t2a* cassette was knocked into the N-terminus of the locus because of different C-terminal splice variants. The self-cleaving T2A peptide frees up GFP::H2B, which will be transported to the nucleus, thereby facilitating cell identification. For *unc-46*, *unc-47*, *tph-1*, *bas-1*, *tbc-1*, *cat-1*, *cat-2*, *snf-3*, and *oct-1*, we knocked-in a *sl2::gfp::h2b* cassette at the C-terminus of the locus. Both types of reporter cassettes should capture posttranscriptional, 3’UTR-mediated regulation of each locus, e.g. by miRNAs and RNA-binding proteins (not captured by CeNGEN scRNA data). Since the reporter is targeted to the nucleus, this strategy circumvents shortcomings associated with interpreting antibody staining patterns or dealing with too densely packed cytosolic signals. For *mod-5*, we analyzed an existing reporter allele (Maicas *et al*. 2021). For all our neuronal cell identification, we utilized the neuronal landmark strain NeuroPAL (Tekieli *et al*. 2021; Yemini *et al*. 2021).

### Expression of a reporter allele of *eat-4/VGLUT*, a marker for glutamatergic identity, in the hermaphrodite

37 of the 38 previously reported neuron classes that express an *eat-4* fosmid-based reporter (Serrano-Saiz *et al*. 2013) showed *eat-4* transcripts in the CeNGEN scRNA atlas (Taylor *et al*. 2021) at all 4 thresholds of stringency, and 1/38 (PVD neuron) showed it in 3 out of the 4 threshold levels (**Fig. 1B**, **Table S1**). However, scRNA transcripts were detected at all 4 threshold levels in three additional neuron classes, RIC, PVN, and DVA, for which no previous reporter data provided support. In a recent publication, we had already described that the *eat-4* reporter allele *syb4257* is expressed in RIC (Reilly *et al*. 2022)(confirmed in **Fig. 4A**). We now also confirm expression of this reporter allele, albeit at low levels, in DVA and PVN (**Fig. 4B**, **Table S2**).

**Figure 3.**
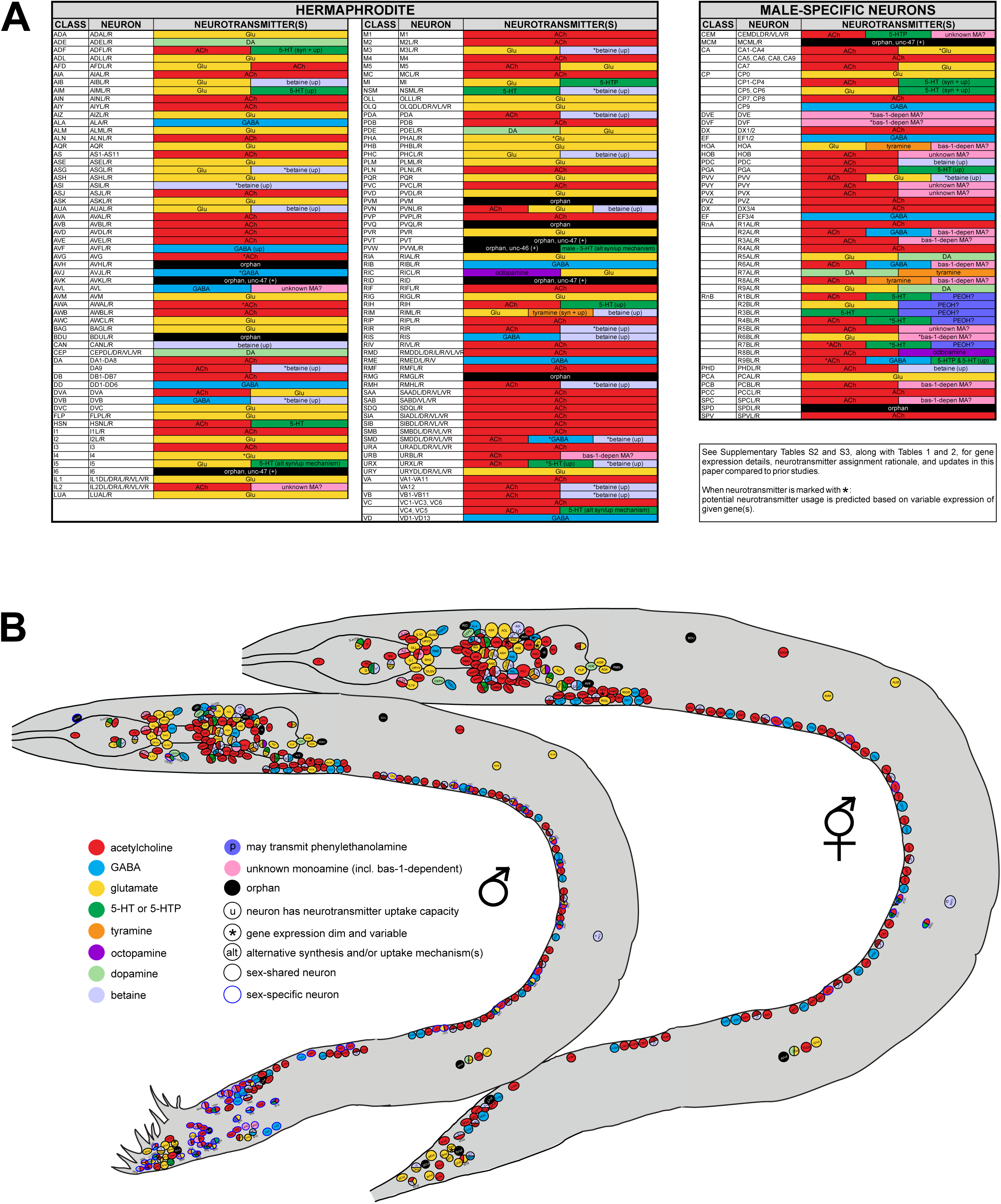
Summary of neurotransmitter usage and atlases. Refer to **Table 1**, **Table 2**, and **Supplementary Tables S2** and **S3** for individual gene expression, rationale for neurotransmitter assignments, and more detailed notes. **(A)** ACh = acetylcholine; Glu = glutamate; DA = dopamine; 5-HT = 5-hydroxytryptamine, or serotonin; 5-HTP = 5-hydroxytryptophan; PEOH? = the neuron has the potential to use β-hydroxyphenethylamine, or phenylethanolamine; bas-1-depen-unknown MA? = the neuron has the potential to use *bas-1*-dependent unknown monoamines (histamine, tryptamine, PEA; also see **Fig. S1**); unknown MA? = the neuron has the potential to use non-canonical monoamines; (up) = neurotransmitter uptake; (syn) = neurotransmitter synthesis; * = dim and variable expression of respective identity gene(s) is detected. Variability could be due to one of the following reasons: (1) the gene is expressed in some but not all animals; (2) the gene is expressed in every animal but the level of expression is below detection threshold in some. Variability is detected only at low fluorescent intensity; at higher intensities, expression remains consistent. Results for anti-GABA staining in SMD and anti-5-HT staining in VC4, VC5, I5, URX, and PVW (male) are variable based on previous reports (see text for citations). **(B)** Information from **A** shown in the context of neuron positions in worm schematics. Note "unknown monoamine" here includes both "*bas-1*-dependent-unknown MA" and "unknown MA" in **A**. Neurons marked with "u" can uptake given neurotransmitters but not exclusively; some may also synthesize them, e.g., ADF can both synthesize and uptake 5-HT.

**Figure 4.**
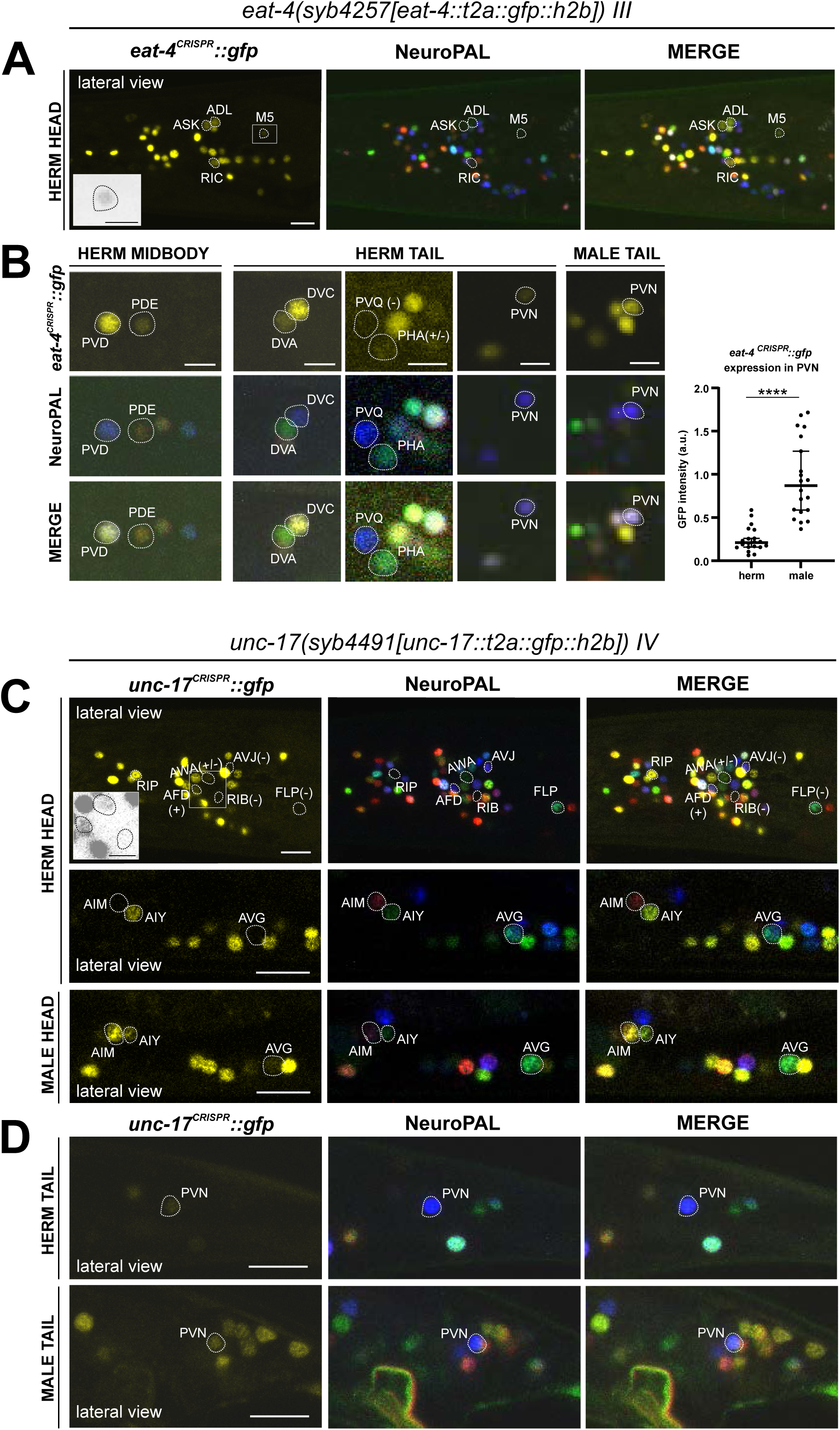
Expression of *eat-4/VGLUT* and *unc-17/VAChT* reporter alleles in the adult hermaphrodite. Neuronal expression of *eat-4(syb4257)* and *unc-17(syb4491)* was characterized with landmark strain NeuroPAL (*otIs696* and *otIs669,* respectively). Only selected neurons are shown for illustrating updates from previous reports. See **Table S2** for a complete list of neurons. **(A)** Dim expression of *eat-4(syb4257)* in head neurons ASK and ADL is consistent with previous fosmid-based reporter expression. RIC expression is consistent with previous observation using the same reporter allele (Reilly *et al*. 2022). In addition, dim expression is detected in pharyngeal neuron M5 (also in grayscale inset), previously not detected with *eat-4* GFP fosmid-based reporter (*otIs388*) but visible with *eat-4* mCherry fosmid-based reporter (*otIs518*). **(B)** Previously uncharacterized *eat-4* expression in PDE and DVA neurons is detected with the *eat-4(syb4257)* reporter allele. Variable expression in PHA is also occasionally detected. No expression is detected in PVQ. Expression in PVN is detected in both sexes but at a much higher level in the male. **(C)** In the head, prominent expression of *unc-17(syb4491)* in RIP and dim expression in AWA and AFD neurons are detected. There is no visible expression in RIB, FLP, or AVJ. Consistent with previous reports, AIM expresses *unc-17* only in males and not hermaphrodites. In addition, very dim expression of AVG can be detected occasionally in hermaphrodites (representative image showing an animal with no visible expression) and slightly stronger in males (representative image showing an animal with visible expression). Inset, grayscale image showing dim expression for AWA and AFD and no expression for RIB. **(D)** In the tail, PVN expresses *unc-17(syb4491)* in both sexes, consistent with previous reports. Scale bars, 10 μm in color images in A, C, and D; 5 μm in B and all grayscale images. Quantification in B is done by normalizing fluorescent intensity of *eat-4* GFP to that of the blue channel in the NeuroPAL background. Statistics, Mann-Whitney test.

Another neuron found to have some *eat-4* transcripts, but only with the two lower threshold sets, is the I6 pharyngeal neuron. Consistent with our previous fosmid-based reporter data, we detected no I6 expression with our *eat-4(syb4257)* reporter allele. The *eat-4* reporter allele also shows expression in the pharyngeal neuron M5, albeit very weakly (**Fig. 4A**, **Table S2**), consistent with CeNGEN scRNA data. Weak expression of the *eat-4* fosmid-based reporter in ASK and ADL remained weak, but clearly detectable with the *eat-4(syb4257)* reporter allele (**Fig. 4A**, **Table S2**). Extremely dim expression in PHA can be occasionally detected. Whereas the PVQ neuron class display *eat-4* scRNA transcripts and was reported to show very dim *eat-4* fosmid-based reporter expression, we detected no expression of the *eat-4(syb4257)* reporter allele in PVQ neurons (**Fig. 4B**, **Table S2**). We also did not detect expression of *eat-4(syb4257)* in the GABAergic AVL and DVB neurons, in which a recent report describes expression using an *eat-4* promoter fusion (Li *et al*. 2023). An absence of *eat-4(syb4257)* expression in AVL and DVB is also consistent with the absence of scRNA transcripts in these neurons.

A few neurons were found to express *eat-4* transcripts by the CeNGEN atlas, but only with lower threshold levels, including, for example, the RMD, PVM, and I4 neurons (**Fig. 1B, Table S1**). We failed to detect reporter allele expression in RMD or PVM neurons, but occasionally observed very dim expression in I4. Lastly, we identified a novel site of *eat-4* expression in the dopaminergic PDE neuron (**Fig. 4B**, **Table S2**). While such expression was neither detected with previous reporters nor scRNA transcripts, we detected it very consistently but at relatively low levels.

### Expression of a reporter allele of *unc-17/VAChT*, a marker for cholinergic identity, in the hermaphrodite

41 of previously described 52 neuron classes that show *unc-17* fosmid-based reporter expression (Pereira *et al*. 2015) showed transcripts in the CeNGEN scRNA atlas at 4 out of 4 threshold levels, another 7 neuron classes at 3 out of 4 threshold levels and 1 at the lowest 2 threshold levels (Taylor *et al*. 2021). Only one neuron class, RIP, displayed scRNA levels at all 4 thresholds, but showed no corresponding *unc-17* fosmid-based reporter expression (**Fig. 1B**, **Table S1**). Using the *unc-17(syb4491)* reporter allele (**Fig. 1A**), we confirmed expression in RIP (**Fig. 4C**, **Table S2**). Of the additional neuron classes that show *unc-17* expression at the lower stringency transcript detection levels (**Fig. 1B**, **Table S1**), we were able to detect *unc-17* reporter allele expression only in AWA (**Fig. 4C**, **Table S2**).

Conversely, a few neurons display weak expression with previous multicopy, fosmid-based reporter constructs (RIB, AVG, PVN)(Pereira *et al*. 2015), but show no CeNGEN scRNA support for such expression (Taylor *et al*. 2021). The *unc-17(syb4491)* reporter allele confirmed weak but consistent expression in the PVN neurons as well as variable, borderline expression in AVG (**Fig. 4C,D**). However, we failed to detect *unc-17(syb4491)* reporter allele expression in the RIB neurons.

We detected a novel site of *unc-17* expression, albeit dim, in the glutamatergic AFD neurons (**Fig. 4C**, **Table S2**). This expression was not reported with previous fosmid-based reporter or CeNGEN scRNA data. scRNA transcript reads for *cha-1/ChAT*, the ACh-synthesizing choline acetyltransferase, were also detected in AFD and PVN (**Table S1**). Taken together, these observations are consistent with the expectation that although available scRNA data capture the majority of gene expression, it may not necessarily have the required depth to detect lowly expressed genes.

Lastly, another notable observation is the lack of any *unc-17* reporter expression or CeNGEN scRNA transcripts in the interneuron AVJ, but presence of CeNGEN scRNA transcript reads for *cha-1/ChAT* (**Table S1**), which shares exons with the *unc-17/VAChT* locus (Alfonso *et al*. 1994). Although no reporter data is available for *cha-1/ChAT*, such interesting mismatch between available *unc-17* and *cha-1/ChAT* expression data could provide a hint to potential non-vesicular cholinergic transmission in the AVJ neurons in *C. elegans*, potentially akin to reportedly non-vesicular release of acetylcholine in the visual system of *Drosophila* (YANG AND KUNES 2004).

### Expression of reporter alleles for GABAergic pathway genes in the hermaphrodite

#### Expression of *unc-25/GAD*

The most recent analysis of GABAergic neurons identified GABA-synthesizing cells by anti-GABA staining and an SL2-based *unc-25/GAD* reporter allele that monitors expression of the rate-limiting step of GABA synthesis, generated by CRISPR/Cas9 engineering (Gendrel *et al*. 2016). The CeNGEN scRNA atlas shows robust support for these assignments at all 4 threshold levels (**Fig. 1B**, **Table S1**). *unc-25* scRNA signals were detected at several orders of magnitude lower levels in 3 additional neuron classes, but only with the least robust threshold level.

In this study we generated another *unc-25/GAD* reporter allele, using a *t2a::gfp::h2b* cassette (*ot1372*) (**Fig. 2**). This allele showed the same expression pattern as the previously described SL2-based *unc-25(ot867)* reporter allele (**Fig. 5A**, **Table S2**). This includes a lack of expression in a number of neurons that stain with anti-GABA antibodies (SMD, AVA, AVB, AVJ, ALA, and AVF) and GLR glia, corroborating the notion that these neurons take up GABA from other cells (indeed, a subset of those cells do express the GABA uptake reporter SNF-11; (Gendrel *et al*. 2016)).

**Figure 5.**
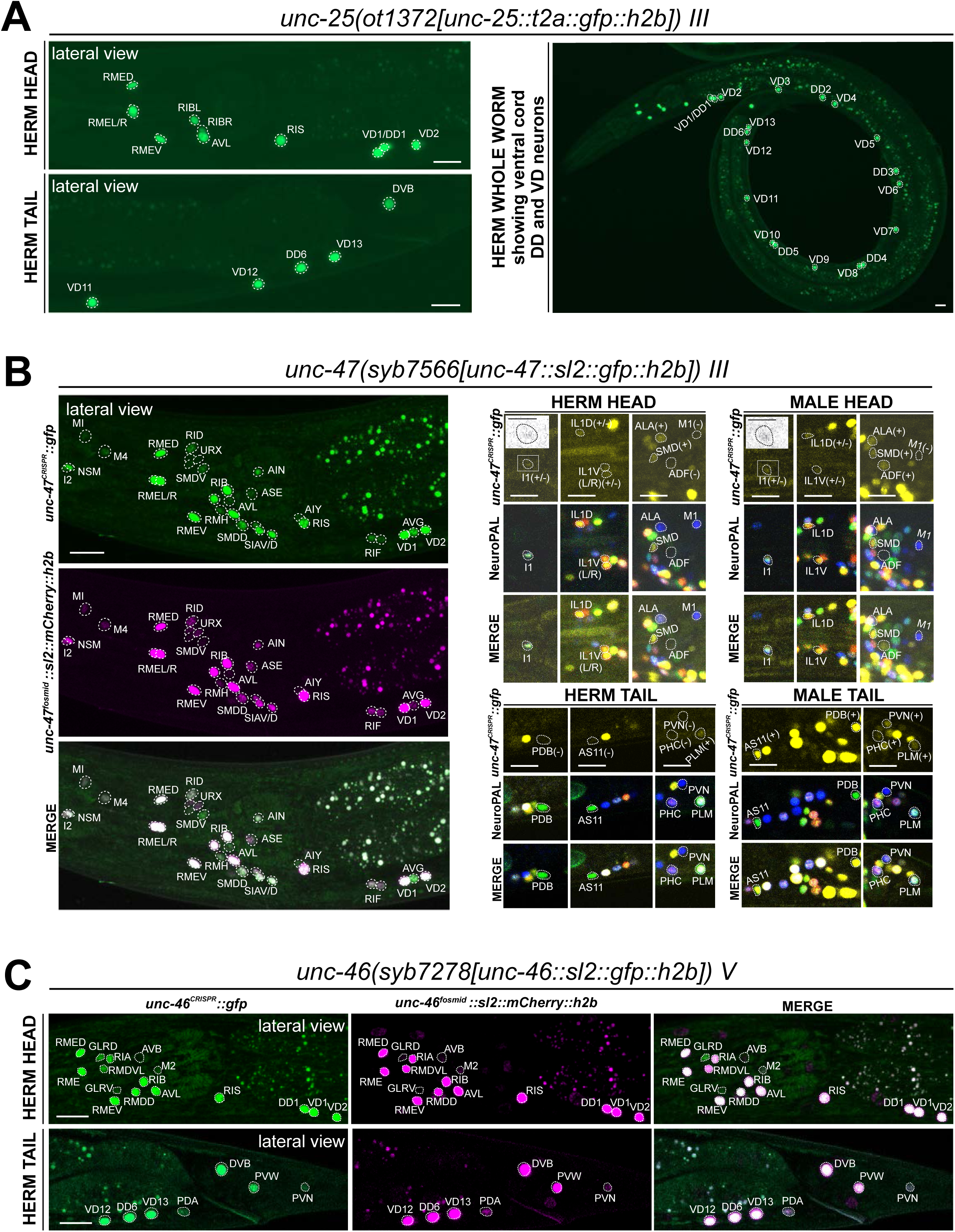
Expression of GABAergic reporter alleles in the adult hermaphrodite. **(A)** Expression of the *unc-25/GAD* reporter allele *ot1372* is detected in the head, ventral nerve cord, and tail neurons. The expression pattern of this new T2A-based reporter allele is similar to that of a previously described SL2-based reporter allele, *unc-25(ot867)*(Gendrel *et al*. 2016). **(B)** Expression of *unc-47/VGAT* reporter allele *syb7566*. **Left**, the expression pattern of the reporter allele largely matches that of a previously described *unc-47* mCherry fosmid-based reporter (*otIs564*) in the head. **Right**, a close-up view for the characterization of the reporter allele expression with landmark strain NeuroPAL (*otIs669*). In the head, consistent with previous reports of the *unc-47* fosmid-based reporter (*otIs564*), dim expression of *unc-47(syb7566)* in SMD, ALA, and very dim and variable expression in IL1 is detected in both sexes, and *unc-47(syb7566)* is expressed in ADF only in the male and not hermaphrodite. In addition, the reporter allele is also expressed at a very dim level in the pharyngeal neuron I1 (also in inset) whereas no expression is detected in M1. In the tail, consistent with previous reports of the fosmid, sexually dimorphic expression of the *unc-47(syb7566)* reporter allele is also detected in PDB, AS11, PVN, and PHC only in the male and not the hermaphrodite. In addition, we also detected very dim expression of PLM in both sexes, confirming potential dim expression of the *unc-47* mCherry fosmid-based reporter that was only readily visible after anti-mCherry staining in the past (Serrano-Saiz *et al*. 2017b). Scale bars, 5 μm for insets and 10 μm for all other images. **(C)** Expression of *unc-46/LAMP* reporter allele *syb7278* is largely similar to that of the previously described *unc-46* mCherry fosmid-based reporter (*otIs568*). We also observed expression of both the reporter allele and fosmid-based reporter in PVW, PVN, and very dimly in PDA. Scale bars, 10 μm.

We carefully examined potential expression in the AMsh glia, which were reported to generate GABA through *unc-25/GAD* (Duan *et al*. 2020; Fernandez-Abascal *et al*. 2022). We did not detect visible *unc-25(ot867)* or *unc-25(ot1372)* reporter allele expression in AMsh, consistent with the failure to directly detect GABA in AMsh through highly sensitive anti-GABA staining (Gendrel *et al*. 2016). Furthermore, since these reporters do not capture an alternatively spiced isoform b.1, we generated another reporter allele, *unc-25(ot1536)*, to specifically target this isoform. However, we did not observe any discernible fluorescent reporter expression from this allele. Hence, it is unlikely that an alternative isoform could contribute to expression in additional cell types.

#### Expression of *unc-47/VGAT*

While promoter-based transgenes for the vesicular transporter for GABA, *unc-47/VGAT*, had shown expression that precisely match that of *unc-25/GAD* (Eastman *et al*. 1999), we had noted in our previous analysis of the GABA system that a fosmid-based reporter showed much broader expression in many additional neuron classes that showed no sign of GABA usage (Gendrel *et al*. 2016). In several of these neuron classes both the fosmid-based reporter and the CeNGEN scRNA data indicate very robust expression (e.g. AIN, SIA, SDQ), while in many others scRNA transcripts are only evident at looser thresholds and, correspondingly, fosmid-based reporter expression in these cells is often weak (**Table S1**) (Gendrel *et al*. 2016). To investigate this matter further, we CRISPR/Cas9-engineered a *gfp-*based reporter allele for *unc-47*, *syb7566*, and first crossed it with an mCherry-based *unc-47* fosmid-based reporter (*otIs564*) as a first-pass assessment for any obvious overlaps and mismatch of expression patterns between the two (**Fig. 5B**, left side panels). The vast majority of neurons exhibited overlapping expression between *syb7566* and *otIs564*. There were also many notable similarities in the robustness of expression of the fosmid-based reporter and the reporter allele (**Table S1**). In a few cases where the fosmid-based reporter expression was so dim that it is only detectable via antibody staining against its fluorophore (mCherry) (Gendrel *et al*. 2016; Serrano-Saiz *et al*. 2017b), the reporter allele expression was readily visible (**Table S1**).

The very few mismatches of expression of the fosmid-based reporter and the reporter allele included the pharyngeal neuron M1, which expresses no visible *unc-47(syb7566)* reporter allele but weak fosmid-based reporter expression, and the pharyngeal neuron I1, which expresses dim *syb7566* but no fosmid-based reporter (**Fig. 5B**, right side panels). Similarly, AVJ shows very dim and variable *unc-47(syb7566)* reporter allele expression but no fosmid-based reporter expression. Since AVJ stains with anti-GABA antibodies (Gendrel *et al*. 2016), this neuron classifies as an uptake and recycling neuron. Other neurons previously shown to stain with anti-GABA antibodies and to express the *unc-47* fosmid-based reporter (ALA and SMD)(Gendrel *et al*. 2016) still show expression of the *unc-47* reporter allele. In contrast, the anti-GABA-positive AVA and AVB neurons express no *unc-47* and therefore possibly operate solely in GABA clearance (see Discussion).

In conclusion, while the reporter allele of *unc-47/VGAT,* in conjunction with CeNGEN scRNA data, corroborates the notion that *unc-47/VGAT* is expressed in all GABA-synthesizing and most GABA uptake neurons, there is a substantial number of *unc-47-*positive neurons that do not show any evidence of GABA presence. This suggests that UNC-47/VGAT may transport another unidentified neurotransmitter (see Discussion)(Gendrel *et al*. 2016).

#### Expression of *unc-46/LAMP*

In all GABAergic neurons, the UNC-47/VGAT protein requires the LAMP-like protein UNC-46 for proper localization (Schuske *et al*. 2007). A previously analyzed fosmid-based reporter confirmed *unc-46/LAMP* expression in all “classic” GABAergic neurons (i.e. anti-GABA and *unc-25/GAD-*positive neurons), but also showed robust expression in GABA- and *unc-47*-negative neurons, such as RMD (Gendrel *et al*. 2016). This non-GABAergic neuron expression is confirmed by CeNGEN scRNA data (Taylor *et al*. 2021)(**Table S1**). We generated an *unc-46/LAMP* reporter allele, *syb7278*, and found its expression to be largely similar to that of the fosmid-based reporter and to the scRNA data (**Fig. 5C, Table S1**), therefore corroborating the non-GABAergic neuron expression of *unc-46/LAMP.* We also detected previously unreported expression in the PVW and PVN neurons in both the reporter allele and fosmid-based reporter (**Fig. 5C**), thereby further corroborating CeNGEN data. In addition, we also detected very dim expression in PDA (**Fig. 5C**), which shows no scRNA transcript reads (**Table S1**). With one exception (pharyngeal M2 neuron class), the sites of non-GABAergic neuron expression of *unc-46/LAMP* expression do not show any overlap with the sites of *unc-47/VGAT* expression, indicating that these two proteins have functions independent of each other.

### Expression of reporter alleles for serotonin biosynthetic enzymes, *tph-1/TPH* and *bas-1/AAAD,* in the hermaphrodite

*tph-1/TPH* and *bas-1/AAAD* code for enzymes required for serotonin (5-HT) synthesis (**Fig. 1A**). scRNA transcripts for *tph-1* and *bas-1* are detected in previously defined serotonergic neurons at all 4 threshold levels (HSN, NSM, ADF) (**Fig. 1**, **Table S1**). In addition to these well characterized sites of expression, several of the individual genes show scRNA-based transcripts in a few additional cells: *tph-1* at all 4 threshold levels in AFD and MI. Neither of these cells display scRNA transcripts for *bas-1/AAAD,* the enzyme that metabolizes the TPH-1 product 5-HTP into 5-HT (**Fig. 1A**). To further investigate these observations, we generated reporter alleles for both *tph-1* and *bas-1* (**Fig. 2**). Expression of the *tph-1* reporter allele *syb6451* confirmed expression in the previously well-described neurons that stained positive for 5-HT, namely NSM, HSN, and ADF, matching CeNGEN data. While expression in AFD (seen at all 4 threshold levels in the CeNGEN scRNA atlas) could not be confirmed with the reporter allele, expression in the pharyngeal MI neurons could be confirmed (**Fig. 6A**, **Table S2**).

**Figure 6.**
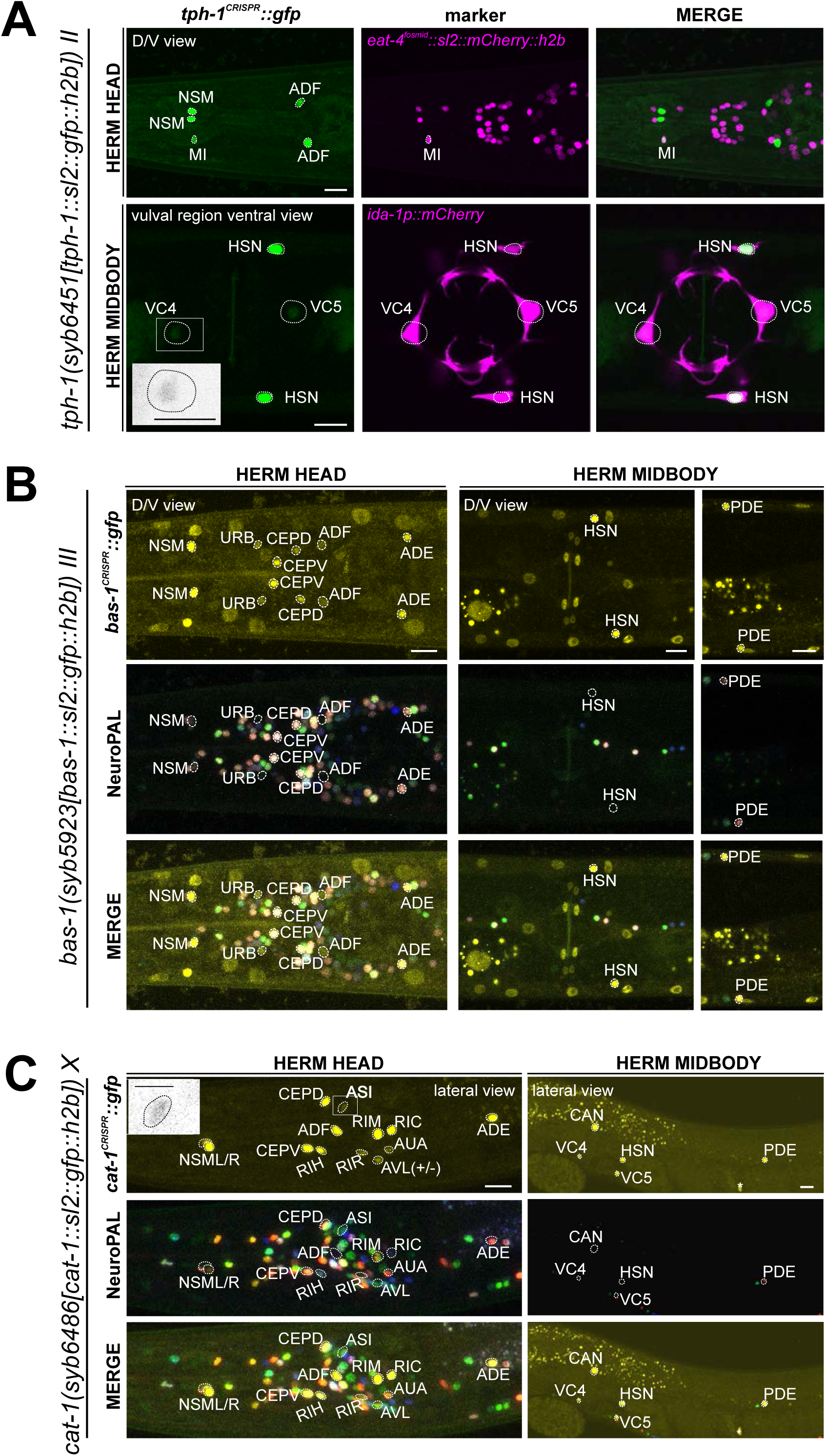
Expression of *tph-1/TPH*, *bas-1/AAAD*, *and cat-1/VMAT* reporter alleles in the adult hermaphrodite. **(A)** Dorsoventral view of a hermaphrodite head and midbody expressing *tph-1(syb6451). tph-1* expression is detected robustly in the MI neuron and dimly and variably in VC4 & VC5. Neuron identities for MI and VC4 & VC5 were validated using *otIs518[eat-4(fosmid)::sl2::mCherry::h2b]* and *vsls269[ida-1::mCherry]*, respectively, as landmarks. Inset, grayscale image highlighting dim expression in VC4. **(B)** Neuronal expression of *bas-1(syb5923)* characterized with the landmark NeuroPAL (*otIs669*) strain in the head and midbody regions of young adult hermaphrodites. Dorsoventral view of the adult head shows *bas-1/AAAD* expression in left-right neuron pairs, including previously reported expression in NSM, CEP, ADF, and ADE (HARE AND LOER 2004). Additionally, we observed previously unreported expression in the URB neurons. Non-neuronal *bas-1/AAAD* expression is detected in other non-neuronal cell types as reported previously (Yu *et al*. 2023) (also see Fig. 14**, S3**). **(C)** Lateral views of young adult hermaphrodite head and midbody expressing *cat-1*/*VMAT* (*syb6486*). Previously unreported *cat-1/VMAT* expression is seen in RIR, CAN, AUA, ASI (also in inset), and variably, ALA. Non-neuronal expression of *cat-1/VMAT* is detected in a single midbody cell in the gonad (also see **Fig. S3**), marked with an asterisk. Scale bars, 10 μm for all color images; 5 μm for the inset in grayscale.

We detected co-expression of the *bas-1* reporter allele, *syb5923*, with *tph-1(syb6451)* in NSM, HSN, and ADF, in accordance with previous reporter and scRNA data (**Fig. 6B**, **Table S2**). However, *bas-1(syb5923)* is not co-expressed with *tph-1* in MI (**Fig. 6A,B**), nor is there CeNGEN-transcript evidence for *bas-1/AAAD* in MI (**Fig. 1**, **Table S1**). Hence, TPH-1-synthesized 5-HTP in MI is not metabolized into 5-HT, consistent with the lack of 5-HT-antibody staining in MI (Horvitz *et al*. 1982; Sze *et al*. 2000).

We also detected *tph-1(syb6451)* reporter allele expression in the serotonergic VC4 and VC5 neurons (**Fig. 6A**, **Table S2**), consistent with scRNA data (**Fig. 1**, **Table S1**) and previous reporter transgene data (Mondal *et al*. 2018). This suggests that these neurons are capable of producing 5-HTP. However, there is no *bas-1(syb5923)* expression in VC4 or VC5, consistent with previous data showing that serotonin is taken up, but not synthesized by them (Duerr *et al*. 2001) (more below on monoamine uptake; **Table 1, 2**).

**Table 1.** Neurons that uptake monoaminergic neurotransmitters. +: presence of reporter allele expression; -: lack of visible reporter allele expression; +/-: dim and variable expression (variability is only detected when reporter fluorescent intensity is low); m: anti-5-HT staining observed in males; *: sex-specific neurons; **: variable/very dim antibody staining reported in previous publications. See text for citations.

| serotonin uptake |  |  |  |  |
| --- | --- | --- | --- | --- |
|  | neuron | *cat-1* | *mod-5* | *tph-1* |
| anti-5-HT staining (+) | ADF | + | + | + |
|  | AIM | - | + | - |
|  | I5** | +/- | - | - |
|  | NSM | + | + | + |
|  | PVW (m) | - | - | - |
|  | RIH | + | + | - |
|  | URX** | - | +/- | - |
|  | *HSN | + | - | + |
|  | *VC4-5** | + | - | +/- |
|  | *CEM** | - | + | + |
|  | *CP1-6 | + | + | + |
|  | *PGA | + | + | - |
|  | *R1B | + | - | + |
|  | *R3B | + | + | + |
|  | *R9B | + | + | + |

| tyramine uptake |  |  |
| --- | --- | --- |
| neuron | *cat-1* | *oct-1* |
| RIM | + | + |

| betaine uptake |  |  |
| --- | --- | --- |
| neuron | *cat-1* | *snf-3* |
| AUA | + | + |
| CAN | + | + |
| NSM | + | +/- |
| RIM | + | + |
| RIR | + | +/- |
| ASI | + | +/- |
| M3 | - | +/- |
| AIB | - | + |
| DVB | - | +/- |
| SMD | - | +/- |
| RIS | - | + |
| URX | - | +/- |
| PDA | - | +/- |
| ASG | - | +/- |
| DA9 | - | +/- |
| VB1-11 | - | +/- |
| PHC | - | + |
| PVN | - | + |
| VA12 | - | +/- |
| RMH | - | +/- |
| *PDC | + | + |
| *PHD | - | + |
| *PVV | - | +/- |

As expected from the role of *bas-1/AAAD* in dopamine synthesis (HARE AND LOER 2004), *bas-1(syb5923)* is also expressed in dopaminergic neurons PDE, CEP, and ADE. In addition, it is also expressed weakly in URB, consistent with scRNA data. We did not detect visible expression in PVW or PVT, both of which showed very low levels of scRNA transcripts (**Fig. 1**, **Table S1**). Expression of *bas-1/AAAD* in URB may suggest that URB generates a non-canonical monoamine (e.g. tryptamine, phenylethylamine, or histamine), but since URB expresses no vesicular transporter (*cat-1/VMAT*, see below), we consider it unlikely that any such monoamine would be secreted via canonical vesicular synaptic release mechanisms.

### Expression of a reporter allele of *cat-2/TH*, a dopaminergic marker, in the hermaphrodite

The CeNGEN scRNA atlas shows transcripts for the rate-limiting enzyme of dopamine synthesis encoded by *cat-2/TH* (**Fig. 1B**, **Table S1**) at all 4 threshold levels in all 3 previously described dopaminergic neuron classes in the hermaphrodite, ADE, PDE, and CEP (Sulston *et al*. 1975; Sulston *et al*. 1980; LINTS AND EMMONS 1999). At lower threshold levels, transcripts can also be detected in the OLL neurons. A CRISPR/Cas9-engineered reporter allele for *cat-2/TH*, *syb8255*, confirmed expression in ADE, PDE and CEP in adult hermaphrodites (**Fig. 7A**, **Table S2**). As expected and described above, all three neuron classes also expressed *bas-1/AAAD* (**Fig. 6B**) and *cat-1/VMAT* (**Fig. 6C**, see below) (**Table S2**). We did not detect visible expression of *cat-2(syb8255)* in OLL. The OLL neurons also display no scRNA transcripts (nor reporter allele expression) for *bas-1/AAAD* or *cat-1/VMAT.* No additional sites of expression of *cat-2(syb8255)* were detected in the adult hermaphrodite.

**Figure 7.**
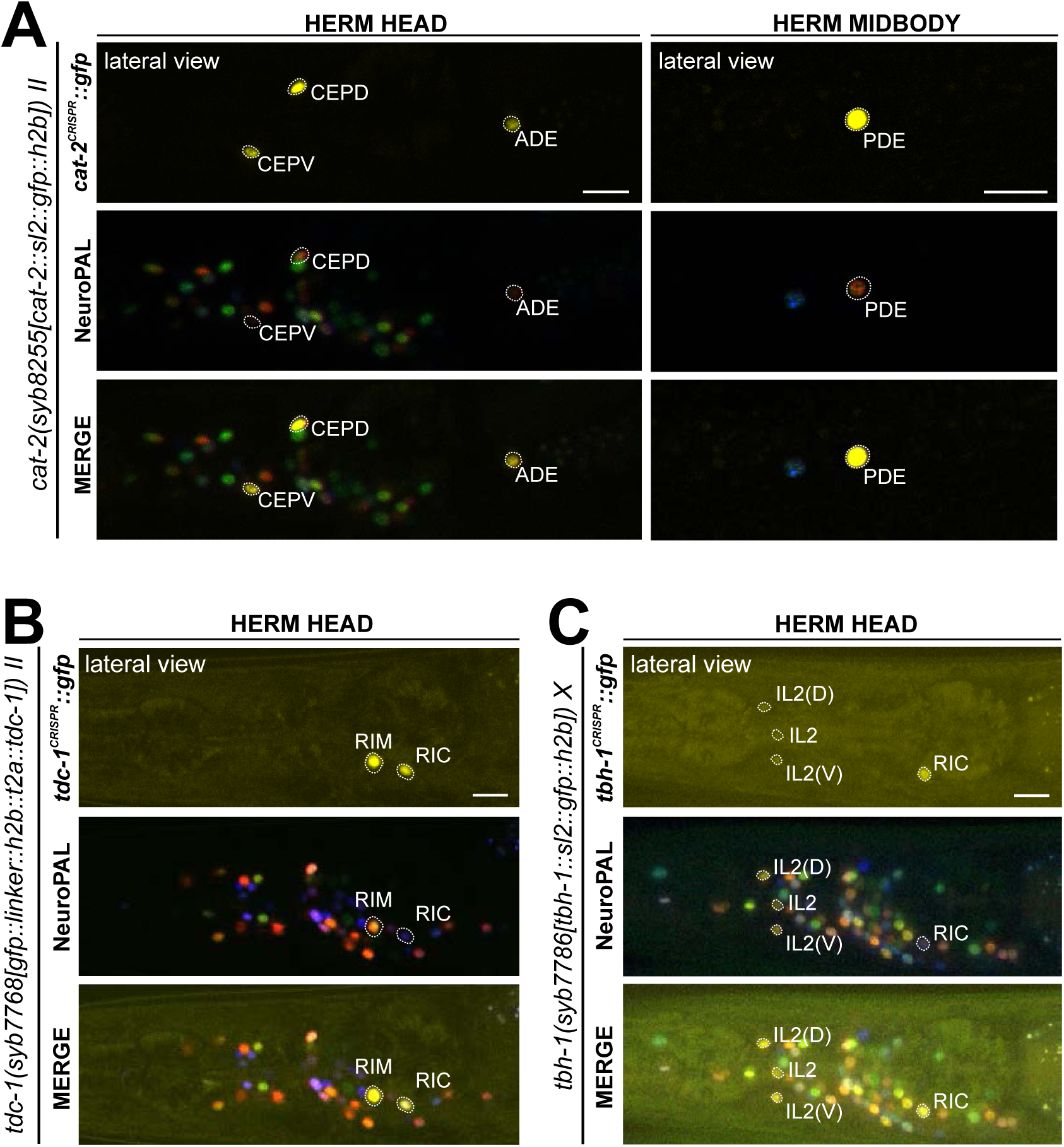
Expression of *cat-2/TH*, *tdc-1/TDC*, and *tbh-1/TBH* reporter alleles in the adult hermaphrodite. Neuronal expression was characterized with landmark strain NeuroPAL (*otIs669*). Lateral views of young adult hermaphrodites expressing reporter alleles for (**A**) *cat-2(syb8255)*, (**B**) *tbh-1(syb7786)*, and (**C**) *tdc-1(syb7768)*. (**A**) *cat-2/TH* expression in CEPs, ADE, and PDE match previously reported dopamine straining expression (Sulston *et al*. 1975). **(B)** and (**C**) Head areas are shown, no neuronal expression was detected in other areas. *tdc-1* expression matches previous analysis (Alkema *et al*. 2005). We detected previously unreported expression of *tbh-1* in all six IL2 neurons at low levels. Scale bars, 10 μm.

### Expression of reporter alleles of *tdc-1/TDC* and *tbh-1/TBH*, markers for tyraminergic and octopaminergic neurons, in the hermaphrodite

The invertebrate analogs of adrenaline and noradrenaline, tyramine and octopamine, are generated by *tdc-1* and *tbh-1* (**Fig. 1A**)(Alkema *et al*. 2005). Previous work had identified expression of *tdc-1* in the hermaphrodite RIM and RIC neurons and *tbh-1* in the RIC neurons (Alkema *et al*. 2005). Transcripts in the CeNGEN atlas match those sites of expression for both *tdc-1* (scRNA at 4 threshold levels in RIM and RIC neurons) and *tbh-1* (scRNA at 4 threshold levels in RIC neurons) (**Fig. 1B, Table S1**). Much lower transcript levels are present in a few additional, non-overlapping neurons (**Fig. 1B**). CRISPR/Cas9-engineered reporter alleles confirmed *tdc-1* expression in RIM and RIC and *tbh-1* expression in RIC (**Fig. 7B,C**, **Table S2**). In addition, we also detected dim expression of *tbh-1(syb7786)* in all six IL2 neurons, corroborating scRNA transcript data (**Fig. 7C**, **Table S2**). However, IL2 neurons do not exhibit expression of the reporter allele of *tdc-1,* which acts upstream of *tbh-1* in the octopamine synthesis pathway, or that of *cat-1/VMAT*, the vesicular transporter for octopamine (**Fig. 6C**, see below). Hence, the IL2 neurons are unlikely to produce or use octopamine as neurotransmitter, but they may synthesize another monoaminergic signal (**Table 2**).

**Table 2.**
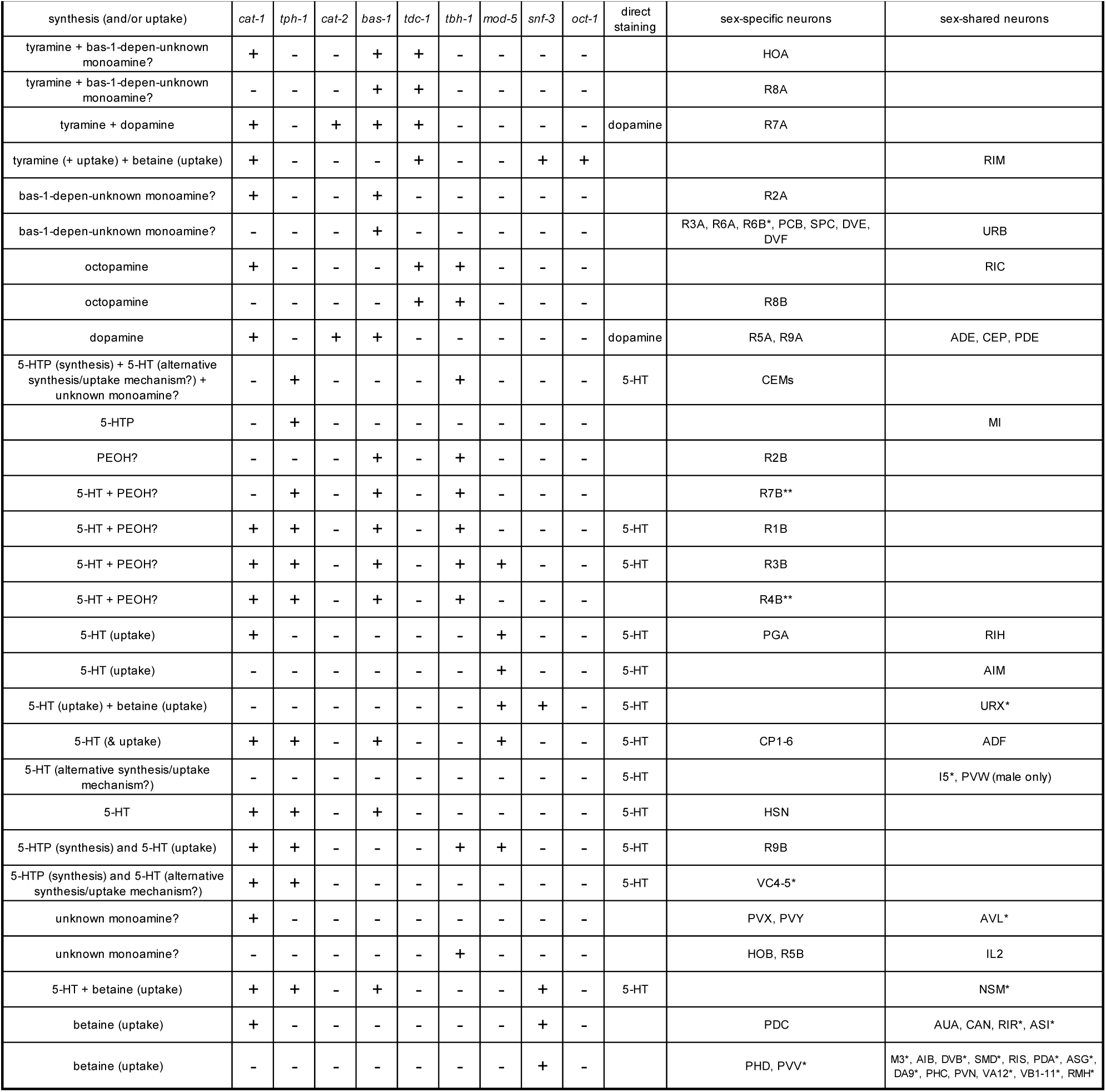
Categories of neuronal expression patterns for monoaminergic neurotransmitter pathway genes. Criteria for monoaminergic neurotransmitter assignment and a summary for neurons with updated identities are presented here. The categories here represent our best assessments based on available data; in every category there is a possibility for the existence of non-canonical synthesis and/or uptake mechanisms that are yet to be discovered. +: presence of reporter allele expression (incl. dim); -: lack of visible reporter allele expression; bas-1-depen-unknown monoamine? = *bas-1*-dependent unknown monoamine (histamine, tryptamine, PEA; see **Fig. S1** and Discussion); unknown monoamine? = potentially non-canonical monoamines; see Discussion and Result sections on specific gene expression patterns; 5-HT = 5-hydroxytryptamine, or serotonin; 5-HTP = 5-hydroxytryptophan; PEOH = β-hydroxyphenethylamine, or phenylethanolamine; *: The expression of *tph-1* in VC4-5, *bas-1* in R4B and R6B, *cat-1* in AVL, and *snf-3* in NSM, RIR, ASI, URX, M3, DVB, SMD, PDA, ASG, DA9, VA12, VB1-11, RMH, and PVV is dim and variable (this study; variability is only detected when reporter fluorescent intensity is low); anti-5-HT staining in VC4, VC5, I5, URX, and PVW (male) is variable in previous reports (see text for citations). ** indicates that R4B and R7B express 5-HT synthesis machinery (*tph-1* and *bas-1*), but do not stain with 5-HT antibodies.

### Expression of a reporter allele of *cat-1/VMAT,* a marker for monoaminergic identity, in the hermaphrodite

As the vesicular monoamine transporter, *cat-1/VMAT* is expected to be expressed in all neurons that synthesize serotonin, dopamine, tyramine, and octopamine (**Fig. 1A**). Both scRNA data and a CRISPR/Cas9-engineered reporter allele, *syb6486*, confirm expression in all these cells (**Fig. 6C**, **Table S2**). In addition, based on antibody staining and previous fosmid-based reporters, *cat-1/VMAT* is known to be expressed in neurons that are serotonin-positive (VC4, VC5, and RIH) (Duerr *et al*. 1999; Duerr *et al*. 2001; Serrano-Saiz *et al*. 2017b). Again, both scRNA data, as well as a CRISPR/Cas9-engineered reporter allele, *syb6486*, confirm expression in these cells (**Fig. 6C**, **Table S2**).

In addition to these canonical monoaminergic neurons, the CeNGEN scRNA data shows *cat-1/VMAT* expression at all 4 threshold levels in RIR, CAN, AVM and, at much lower threshold, 8 additional neuron classes (**Fig. 1B**, **Table S1**). Our *cat-1/VMAT* reporter allele, *syb6486,* corroborates expression in RIR and CAN, but not in AVM (**Fig. 6C**, **Table S2**). We also observed expression of the *cat-1* reporter allele in two of the neuron classes with scRNA transcripts at the lowest threshold level, ASI and variably, AVL (**Fig. 6C, Table S1**). Interestingly, AVL does not express any other monoaminergic pathway genes (**Table S2**), therefore it may be transporting a new amine yet to be discovered. This scenario also applies for two male-specific neurons (more below). As previously mentioned, we detected no *cat-1/VMAT* expression in the *tph-1-*positive MI or the *cat-2/TH-*positive OLL neurons.

The *cat-1/VMAT* reporter allele revealed expression in an additional neuron class, the AUA neuron pair (**Fig. 6C**, **Table S2**). Expression in this neuron is not detected in scRNA data; such expression may be consistent with previous CAT-1/VMAT antibody staining data (Duerr *et al*. 1999). These authors found the same expression pattern as we detected with *cat-1/VMAT* reporter allele, except for the AIM neuron, which Duerr *et al*. identified as CAT-1/VMAT antibody-staining positive. However, neither our reporter allele, nor a fosmid-based *cat-1/VMAT* reporter, nor scRNA data showed expression in AIM, and we therefore think that the neurons identified by Duerr *et al* as AIM may have been the AUA neurons instead (Serrano-Saiz *et al*. 2017b). Additionally, a *cat-1*-positive neuron pair in the ventral ganglion, unidentified but mentioned by Duerr and colleagues (Duerr *et al*. 1999), is likely the tyraminergic RIM neuron pair, based on our reporter allele and CeNGEN scRNA data.

### Expression of reporter alleles of monoamine uptake transporters in the hermaphrodite

In addition to or in lieu of synthesizing monoamines, neurons can uptake them from their surroundings. To investigate the cellular sites of monoamine uptake in more detail, we analyzed fluorescent protein expression from engineered reporter alleles for the uptake transporters of 5-HT (*mod-5/SERT(vlc47)*), the predicted uptake transporter for tyramine (*oct-1/OCT(syb8870)*), and that for betaine (*snf-3/BGT1(syb7290)*).

#### Serotonin/5-HT uptake

Using a promoter-based transgene and antibody staining, previous work had shown expression of the serotonin uptake transporter *mod-5* in NSM, ADF, RIH, and AIM (Jafari *et al*. 2011; Maicas *et al*. 2021). This matched the observations that RIH and AIM do not synthesize 5-HT (i.e. do not express *tph-1*), but stain positive with a 5-HT antibody (Jafari *et al*. 2011). In *mod-5* mutants or wildtype worms treated with serotonin reuptake inhibitors (such as the SSRI fluoxetine), RIH and AIM lose 5-HT immunoreactivity (Jafari *et al*. 2011). We analyzed a CRISPR-based reporter allele, *mod-5(vlc47)*(MAICAS et al. 2021), and confirmed expression in the four neuron classes NSM, ADF, RIH, and AIM (**Fig. 8**). Because only NSM, ADF, and RIH, but not AIM, express the reporter allele of the monoamine transporter CAT-1/VMAT (**Fig. 6**), we agree with previous studies that AIM likely functions as a serotonin uptake/clearance neuron (**Table 1, 2**; see also Discussion). In addition, we also detected dim expression in the phasmid neuron class PHA and very dim, variable signal in URX (**Fig. 12A,B,E**), consistent with scRNA data (**Table S1**). The results for anti-5-HT-staining from previous reports are variable in a few neurons, possibly due to differences in staining methods (including URX, I5, VC4, VC5 and PVW (LOER AND KENYON 1993; RAND AND NONET 1997; Duerr *et al*. 1999; Serrano-Saiz *et al*. 2017b). In light of its *mod-5* reporter expression, URX may acquire 5-HT via *mod-5*, akin to AIM (**Table 1, 2**).

**Figure 8.**
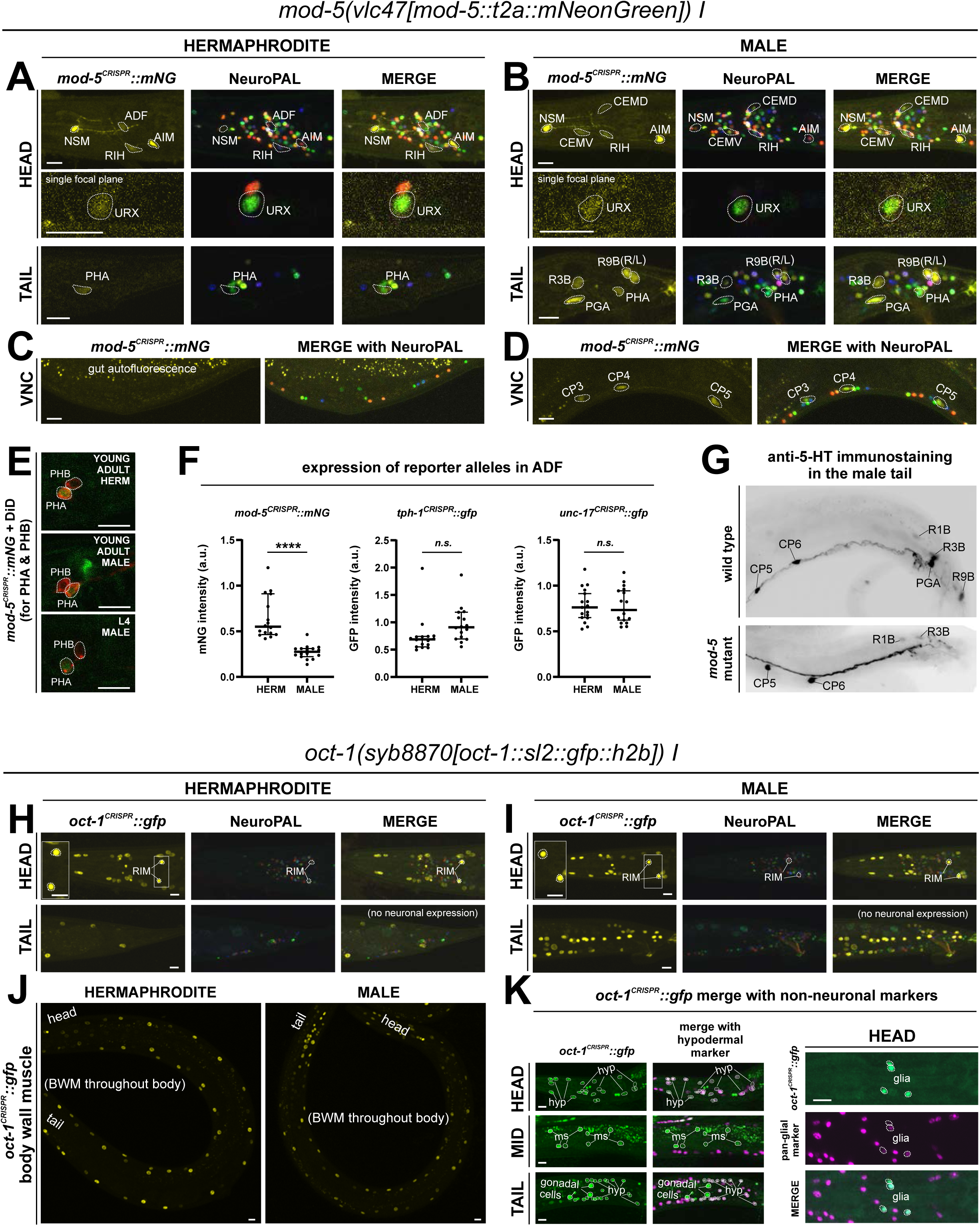
Expression of *mod-5/SERT* and *oct-1/OCT* reporter alleles in adult animals. Neuronal expression was characterized with landmark strain NeuroPAL (*otIs669*) and DiD-filling. (**A,C**) In adult hermaphrodites, *mod-5(vlc47)* is expressed in sex-shared neurons NSM, ADF, RIH, AIM, consistent with previous reports(Jafari *et al*. 2011; Maicas *et al*. 2021). In addition, we also observed expression in the phasmid neuron PHA and dim and variable expression in URX. There is no visible expression in the ventral nerve cord (VNC). (**B,D**) In adult males, *mod-5(vlc47)* is visibly expressed in NSM, RIH, AIM, as well as the male-specific neurons CEM, PGA, R3B, R9B, and CP1 to CP6. Expression in ADF is often not detected (see F). **(E)** DiD-filling confirms *mod-5(vlc47)* expression in phasmid neuron class PHA, and not PHB, in young adults in both sexes (L4 male image is to facilitate neuron ID in adults, because the positions of the two neuron classes can change in males during the L4 to adult transition). **(F)** Expression of *mod-5(vlc47)* in ADF is stronger in hermaphrodites than in males. In comparison, expression is not sexually dimorphic for the reporter alleles of either the 5-HT-synthsizing enzyme *tph-1* or the vesicular acetylcholine transporter *unc-17*. **(G)** In the tail region of wild type males, male-specific neurons PGA, R1B, R3B, and R9B are stained positive for 5-HT. In a *mod-5(n3314)* mutant background, staining is completely lost in PGA (41/41 stained animals) and significantly affected for R9B (completely lost in 31/41 animals and much dimmer in the rest), while it remains in all 41 stained animals for R1B and R3B. The staining for CP1 to CP6 are also not affected in *mod-5* mutant animals (remaining in 41/41 stained animals; image showing CP5 and CP6). (**H,I**) In adult animals, *oct-1(syb8870)* is expressed in the tyraminergic neuron class RIM in both sexes. Expression is not observed in any other neurons. (**J,K**) Outside the nervous system, *oct-1(syb8870)* is expressed in body wall muscle (BWM) throughout the worm (**J**) as well as hypodermal cells and selected head glia (**K**). Expression is also observed in gonadal cells in the male vas deferens (**K**). A pan-glial reporter *otIs870[mir-228p::3xnls::TagRFP]* and a *dpy-7p::mCherry reporter stIs10166 [dpy-7p::his-24::mCherry + unc-119(+)]* were used for glial and hypodermal identification, respectively. Scale bars, 10 μm.

In the hermaphrodite-specific neurons HSN, VC4, and VC5, we did not observe expression of the *mod-5* reporter allele (**Table 1, 2**). Since VC4 and VC5 do not express the complete synthesis pathway for 5-HT, we infer that the anti-5-HT staining in these neurons is a result of alternative 5-HT uptake or synthesis mechanisms. A similar scenario holds for the pharyngeal neuron I5 which was previously reported to stain weakly for 5-HT (Serrano-Saiz *et al*. 2017b).

#### Tyramine uptake

Biochemical studies in vertebrates have shown that the SLC22A1/2/3 (aka OCT-1/2/3) organic anion transporter can uptake monoaminergic neurotransmitters (NIGAM 2018), with SLC22A2 being apparently selective for tyramine (Berry *et al*. 2016). *oct-1* is the ortholog of the OCT subclass of SLC22 family members (Zhu *et al*. 2015), but neither its expression nor function in the nervous system had been previously reported. We tagged the endogenous *oct-1* locus with an *sl2::gfp::h2b* cassette (*syb8870*) and, within the nervous system, observed exclusive expression in the RIM neuron (**Fig. 8H,I**), indicating that RIM is likely capable of uptaking tyramine in addition to synthesizing it via *tdc-1/TDC*. This is consistent with RIM being the only neuron showing *oct-1* scRNA transcripts at all 4 threshold levels in the CeNGEN atlas (**Table S1**).

#### Betaine uptake

Notably, four CAT-1/VMAT- expressing neuron classes, CAN, AUA, RIR, and ASI do not express biosynthetic enzymes for synthesis of the four conventional monoaminergic transmitters known to be employed in *C. elegans* (5-HT, dopamine, octopamine, or tyramine). Hence, these neuron classes might instead uptake some of these transmitters. We considered the putative neurotransmitter betaine as a possible candidate, since CAT-1/VMAT is also able to package betaine (Peden *et al*. 2013; Hardege *et al*. 2022). Betaine is synthesized endogenously, within the nervous system mostly in the *cat-1/VMAT-* positive RIM neuron (Hardege *et al*. 2022), but it is also available in the bacterial diet of *C. elegans* (Peden *et al*. 2013). In vertebrates, dietary betaine is taken up by the betaine transporter BGT1 (i.e. SLC6A12). To test whether *cat-1/VMAT*-positive neurons may acquire betaine via BGT1-mediated uptake, we CRISPR/Cas9-engineered a reporter allele for *snf- 3/BGT1*, *syb7290*. We detected expression in the betaine-synthesizing (and also tyraminergic) RIM neuron (**Fig. 9**, **Table 1, 2**). In addition, *snf-3* is indeed expressed in all the four *cat-1/VMAT-*positive neuron classes that do not synthesize a previously known monoaminergic transmitter (CAN, AUA, and variably, RIR and ASI)(**Fig. 9A,B**). These neurons may therefore take up betaine and synaptically release it via CAT-1/VMAT. The *snf-3(syb7290)* reporter allele is also expressed in the serotonergic neuron NSM (albeit variably) (**Table 1, 2**), thus NSM could also be a betaine uptake neuron. In addition, we also detected *snf-3(syb7290)* expression in several other neurons that do not express *cat-1(syb6486)* (**Table S1**). Expression was also observed in a substantial number of non-neuronal cell types (**Fig. 9E-G**, **Table 2, S1**). These neurons and non-neuronal cells may serve to clear betaine (see Discussion, Neurotransmitter synthesis versus uptake). *snf-3(syb7290)* is not expressed in the inner and outer labial neuron classes as previously suggested (Peden *et al*. 2013); these cells were likely misidentified in the previous study and are in fact inner and outer labial glial cells (as discussed further below).

**Figure 9.**
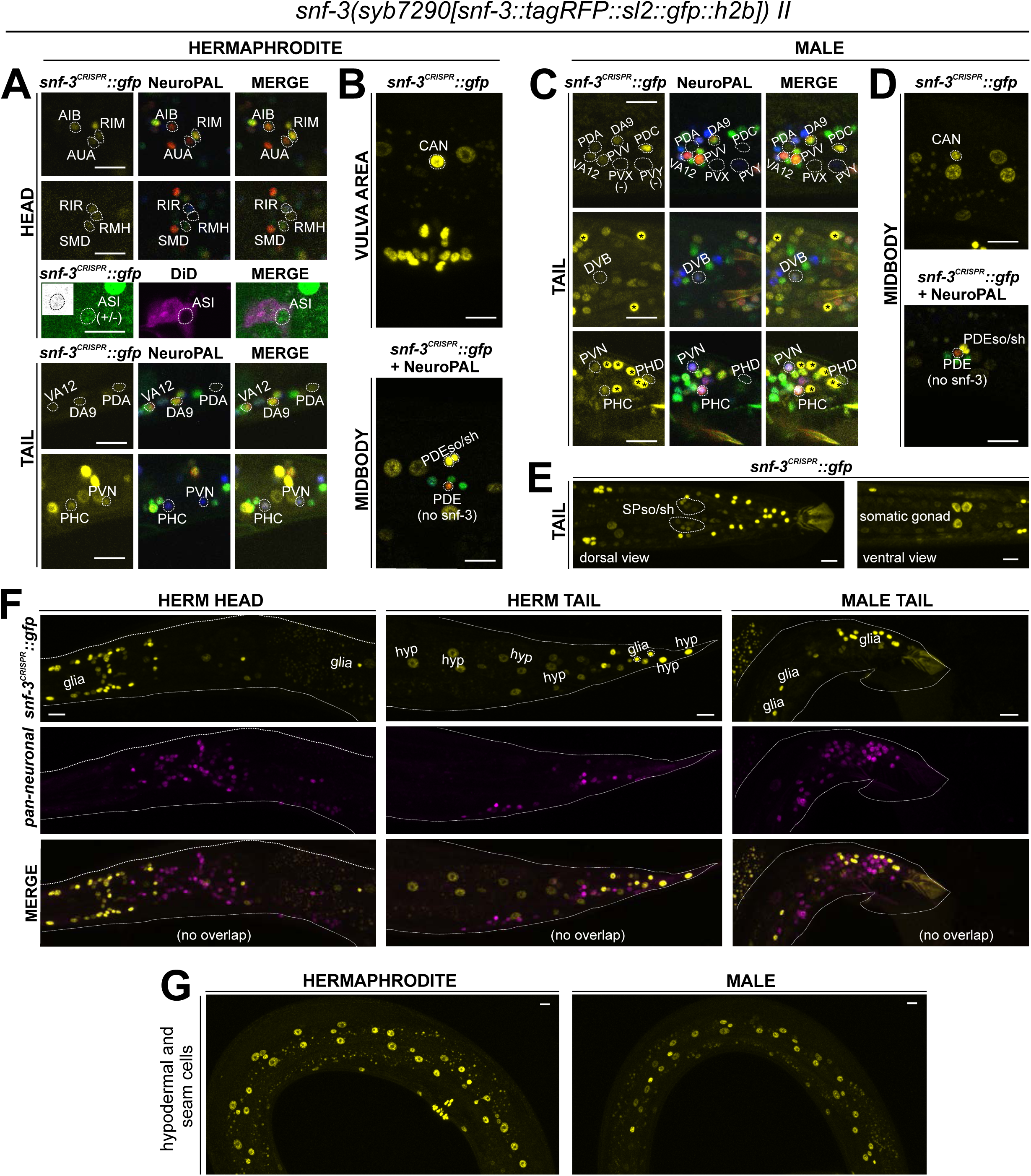
Expression of *snf-3/BGT1/SLC6A12* in adult animals. Neuronal expression was characterized with landmark strain NeuroPAL (*otIs669*) and DiD-filling. (**A,B**) In the adult hermaphrodite, neuronal expression of *snf-3(syb7290)* is detected in *cat-1/VMAT*-positive neurons AUA, CAN, and dimly and variably, RIR and ASI (confirmed with DiD-filling). In addition, it is also expressed in *cat-1/VMAT*-negative neurons AIB, RIM, RMH, SMD, VA12, DA9, PDA, PHC, PVN as labeled, as well as more neurons listed in **Table S1**. In the midbody, expression is not in PDE (dopaminergic, *cat-1*-positive) but in its associated glial cells. It is also detected in multiple vulval support cells (**B**) and some epithelial cells near the somatic gonad. **(C)** In the adult male, in addition to its expression in sex-shared neurons as in hermaphrodites, *snf-3(syb7290)* is also expressed in male-specific neuron class PDC, as well as in PHD and variably in PVV. **(D)** Similarly to its expression in hermaphrodites, *snf-3(syb7290)* is detected in CAN and PDE-associated glial cells, but not PDE neurons, in males. **(E)** In the male tail, *snf-3(syb7290)* is expressed in a number of glial cells including the spicule sockets and/or sheath cells (dorsal view). It is also detected in the somatic gonad (ventral view). **(F)** *snf-3(syb7290)* is broadly expressed in most if not all glia in both sexes. Glial cell type is determined by cell location and the appearance of their nuclei in Normarski. To confirm they are not neurons, a pan-neuronal marker (UPN, or “uber pan-neuronal”, a component in NeuroPAL) is used to determine non-overlapping signals between the two reporters. Head expression in the male is very similar to that in the hermaphrodite and thus not shown. (**G**) *snf-3(syb7290)* is broadly expressed in hypodermal and seam cells in both sexes. Scale bars, 10 μm. Asterisks, non-neuronal expression.

Together with the expression pattern of the uptake transporters, all *cat-1/VMAT*-positive neurons in the hermaphrodite can be matched with an aminergic neurotransmitter. We nevertheless wondered whether another presently unknown monoaminergic transmitter, e.g., histamine or other trace amine, could be synthesized by a previously uncharacterized AAAD enzyme encoded in the *C. elegans* genome, *hdl-1* (**Fig. S1A**)(HARE AND LOER 2004). We CRISPR/Cas9-engineered an *hdl-1* reporter allele, *syb1048*, but detected no expression of this reporter in the animal (**Fig. S1C,D**). Attempts to amplify weak expression signals by insertion of Cre recombinase into the locus failed [*hdl-1(syb4208)*](see Methods). CeNGEN scRNA data also shows no strong transcript expression in the hermaphrodite nervous system and only detected notable expression in sperm (Taylor *et al*. 2021).

### Reporter alleles and NeuroPAL-facilitated neuron class-identification reveal novel expression patterns of neurotransmitters in the male-specific nervous system

No comprehensive scRNA atlas has yet been reported for the nervous system of the male. Based on the expression of fosmid-based reporters, we had previously assembled a neurotransmitter atlas of the *C. elegans* male nervous system in which individual neuron classes are notoriously difficult to identity (Serrano-Saiz *et al*. 2017b). We have since established a NeuroPAL landmark strain that permits more reliable identification of gene expression patterns in both the hermaphrodite and male-specific nervous system (Tekieli *et al*. 2021; Yemini *et al*. 2021). We used NeuroPAL to facilitate the analysis of the expression profiles of our CRISPR/Cas9-engineered reporter alleles in the male, resulting in updated expression profiles for 11 of the 16 reporter alleles analyzed. As in the hermaphrodite, reasons for the updates vary. In addition to the improved accuracy of neuron identification provided by NeuroPAL, in some cases there are true differences of expression patterns between the fosmid-based reporters and reporter alleles. We elaborate on these updates for individual reporter alleles below.

### Expression of reporter alleles of Glu/ACh/GABA markers in the male-specific nervous system

We analyzed *eat-4/VGLUT* (*syb4257*), *unc-17/VAChT* (*syb4491*), *unc-25/GAD* (*ot1372*), and *unc-47/VGAT* (*syb7566*) expression in the male-specific nervous system using NeuroPAL landmark strains (*otIs696* for *eat-4* and *otIs669* for all others). Of all those reporter alleles, *unc-25/GAD* (*ot1372*) was the only one with no updated expression. Specifically, in addition to confirming presence of expression of the *unc-25(ot1372)* reporter allele in CP9, EF1/2, EF3/4, we also confirmed its *lack* of expression in anti-GABA-positive neurons R2A, R6A, and R9B (Gendrel *et al*. 2016; Serrano-Saiz *et al*. 2017b)(**Fig. 11A**, **Table S3**).

In the preanal ganglion, we observed weak expression of *unc-17(syb4491) in* DX3/4 (**Fig. 10B**, **Table S3**), hence assigning previously unknown neurotransmitter identity to these neurons. Related to DX3/4, we also confirmed expression of *unc-17* in DX1/2 in the dorsorectal ganglion, consistent with fosmid-based reporter data (**Table S3**) (Serrano-Saiz *et al*. 2017b). In the lumbar ganglion, we detected novel expression of *unc-17(syb4491)* in 5 pairs of type B ray neurons, namely R1B, R4B, R5B, R7B, and R9B (**Fig. 10B**, **Table S3**). Expression in all these neurons is low, possibly explaining why it is not observed with an *unc-17* fosmid-based reporter (Serrano-Saiz *et al*. 2017b).

**Figure 10.**
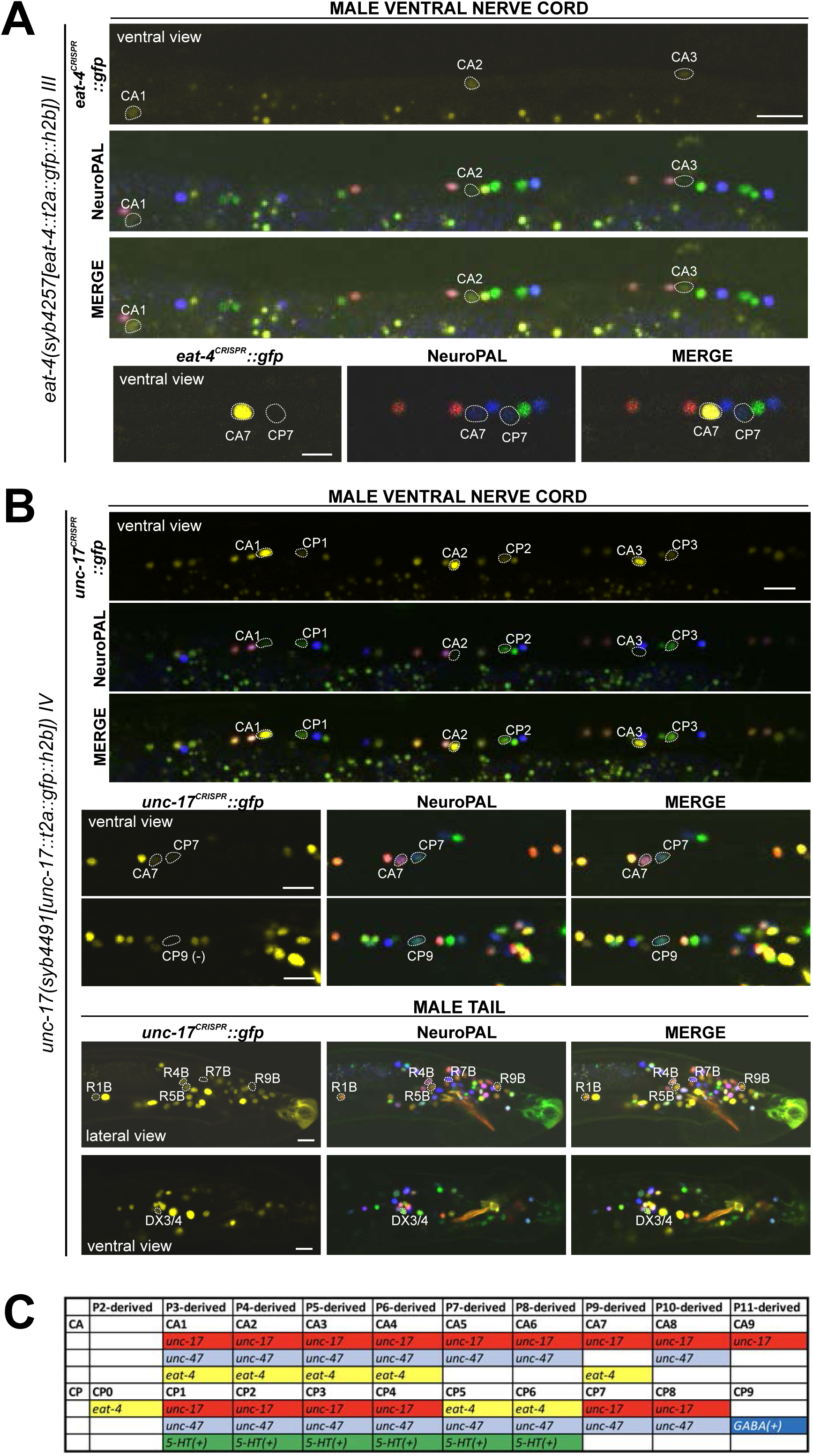
Expression of *eat-4/VGLUT* and *unc-17/VAChT* reporter alleles in the adult male. Neuronal expression of *eat-4(syb4257)* and *unc-17(syb4491)* was characterized with landmark strain NeuroPAL (*otIs696* and *otIs669*, respectively). Only selected neurons are shown to illustrate updates from previous studies. See **Table S3** for a complete list of neurons. **(A)** *eat-4(syb4257)* expression. Top, long panels: CA1, CA2, and CA3 show visible, albeit very dim, novel expression of *eat-4* (also expressed in CA4). Bottom panels: CA7 strongly expresses *eat-4(syb4257)*, whereas CP7 does not. Neuron IDs for these two neurons were previously switched (Serrano-Saiz *et al*. 2017b). **(B)** *unc-17(syb4491)* expression. Top, long panels: ventral view of a male ventral nerve cord showing high levels of expression in CA1, CA2, and CA3 and previously unreported low levels of expression in CP1, CP2, and CP3. Middle panels: low levels of expression in CA7 and CP7. There is no visible expression in CP9. Bottom panels: lateral view of a male tail showing previously unreported dim expression in R1B, R4B, R5B, R7B, and R9B; ventral view of the pre-anal ganglion showing expression in DX3/4. Scale bars, 10 μm. **(C)** The updated neurotransmitter atlas underscores the molecular diversity of the male-specific ventral cord neuron class CA and CP. Based on their expression patterns for neurotransmitter genes, these neurons can be potentially grouped into the following 4 CA and 5 CP sub-classes. See **Table S3** and **Figs. 10-12** for all genes mentioned in the following. CAs: (1) CA1 to CA4: express *unc-17*, *unc-47*, very weak *eat-4*; (2) CA5, 6, 8: express *unc-17* and *unc-47*; (3) CA7: expresses *unc-17*, strong *eat-4*, and no *unc-47*; (4) CA9: expresses weak *unc-17*. CPs: (1) CP0: only expresses weak *eat-4*; (2) CP1 to CP4: express *unc-17*, *unc-47*, *cat-1*, *tph-1*, *bas-1*, and stain for 5-HT; (3) CP5, 6: express *eat-4*, *unc-47*, *cat-1*, *tph-1*, *bas-1*, and stain for 5-HT; (4) CP7, 8: express very weak *unc-17* and *unc-47*; (5) CP9: expresses strong *unc-25* and *unc- 47*, and stain for GABA. The proposed sub-classification for these neuron classes indicates different functions of individual CA and CP neurons; it also provides useful resources for ongoing single-cell sequencing efforts for male-specific neurons.

**Figure 11.**
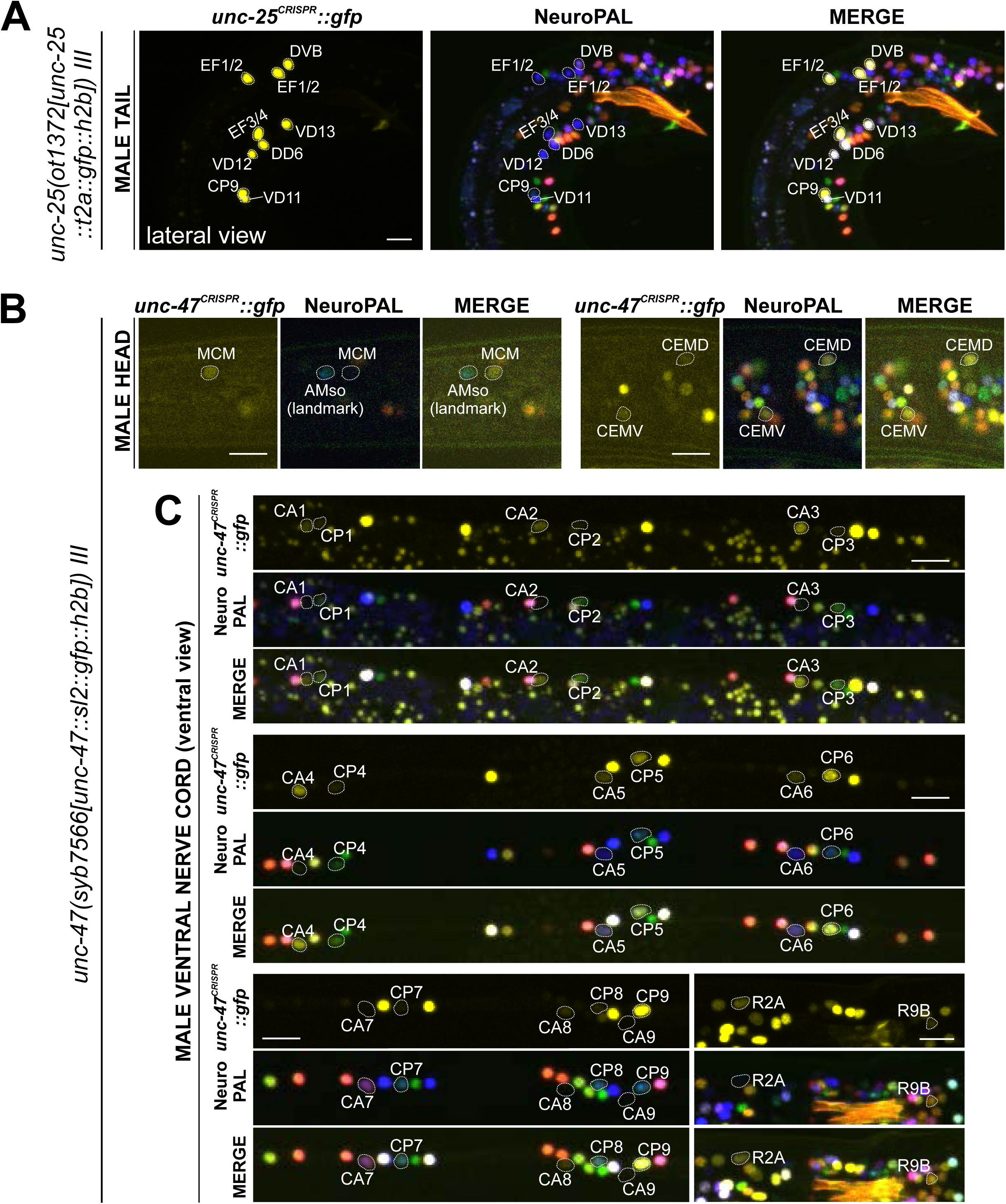
Expression of GABAergic reporter alleles in the adult male. Neuronal expression of *unc-25(ot1372)* and *unc-47(syb7566)* reporter alleles was characterized with landmark strain NeuroPAL (*otIs669*). Only selected neurons are shown to illustrate updates from previous reports. See **Table S3** for a complete list of neurons. **(A)** *unc-25(ot1372)* is expressed in male-specific CP9 and EF neurons as well as a few sex-shared neurons, all consistent with previous reports (Gendrel *et al*. 2016; Serrano-Saiz *et al*. 2017b). **(B)** *unc-47(syb7566)* shows expression in male head neuron classes MCM and CEM, the former previously undetected and the latter consistent with fosmid-based reporter *otIs564*. **(C)** *unc-47(syb7566)* shows expression in a number of ventral cord CA and CP neurons, largely consistent with reported *otIs564* fosmid-based reporter expression except for no visible expression of *syb7566* in CA7 (due to its initial confusion with CP7, described in Fig. 10) and presence of very dim expression in CP7. The *syb7566* reporter allele is also not visible in CA9. Scale bars, 10 μm.

In the ventral nerve cord, we found additional, very weak expression of *eat-4(syb4257)* in CA1 to CA4 (**Fig. 10A, Table S3**), as well as weak expression of *unc-17(syb4491)* in CP1 to CP4 (**Fig. 10B**, **Table S3**), all undetected by previous analysis of fosmid-based reporters (Serrano-Saiz *et al*. 2017b). Conversely, two neurons lack previously reported expression of fosmid-based reporters; CP9 does not show visible *unc-17(syb4491)* expression (**Fig. 10B**) and neither does CA9 show visible expression of *unc-47(syb7566)* expression (**Fig. 11C**). We also realized that the neuron identifications of CA7 and CP7 were previously switched (Serrano-Saiz *et al*. 2017b), due to lack of proper markers for those two neurons. With NeuroPAL, we are now able to clearly distinguish the two and update their classic neurotransmitter reporter expression: CA7 expresses high levels of *eat-4(syb4257)* (**Fig. 10A**, **Table S3**), very low levels of *unc-17(syb4491)* (**Fig. 10B**), and no *unc-47(syb7566)* (**Fig. 10C**); CP7 expresses no *eat-4(syb4257)* (**Fig. 10A**, **Table S3**), very low levels of *unc-17(syb4491)* (**Fig. 8B**), and very low levels of *unc-47(syb7566)* as well (**Fig. 11C**). Taken together, the analysis of reporter alleles reveals a remarkable diversity of CA and CP neurons, summarized in **Fig. 8C**.

In the head, we detected expression of *unc-47(syb7566)* in the male-specific neuron class MCM (**Fig. 11B**, **Table S3**), previously not observed with fosmid-based reporters. Consistent with fosmid-based reporter data, the other male-specific neuron class, CEM, shows expression of *unc-17(syb4491)* (**Table S3**) and *unc-47(syb7566)* (**Fig. 11B**, **Table S3**) reporter alleles.

### Expression of reporter alleles for monoaminergic neurotransmitter pathway genes in the male-specific nervous system

We analyzed the expression of reporter alleles for the following genes involved in monoamine biosynthesis and uptake in the male-specific nervous system: *cat-1/VMAT* (*syb6486*), *tph-1/TPH* (*syb6451*), *cat-2/TH* (*syb8255*), *bas-1/AAAD* (*syb5923*), *tdc-1/TDC* (*syb7768*), *tbh-1/TBH* (*syb7786*), *mod-5/SERT* (*vlc47*), *oct-1/OCT* (*syb8870*), and *snf-3/BGT1* (*syb7290*). As in the hermaphrodite nervous system, we used the NeuroPAL reporter landmark (*otIs669*) for neuron ID (Tekieli *et al*. 2021). We found novel expression patterns in all male-specific ganglia (**Fig. 12, 13**, **Table S3**).

**Figure 12.**
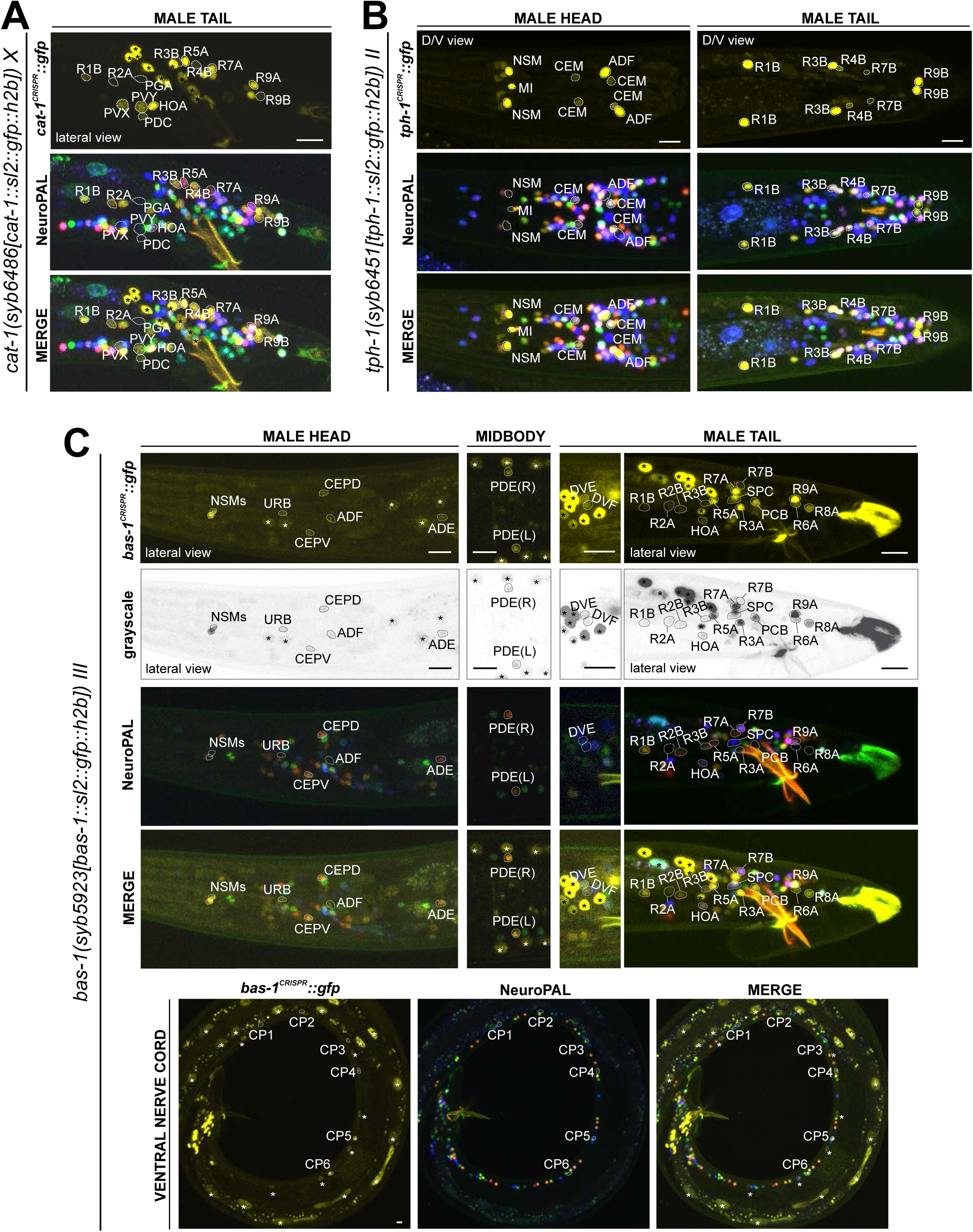
Expression of the *cat-1/VMAT*, *tph-1/TPH*, and *bas-1/AAAD* reporter alleles in the adult male. Neuronal expression was characterized with landmark strain NeuroPAL (*otIs669*). **(A)** Novel *cat-1(syb6486)* expression is seen in male-specific neurons PDC, PVY, PVX, R2A, and R4B. Consistent with previous reports, it is also expressed in HOA, PGA, R5A, R7A, R9A, R1B, and R8B. Its expression in ventral cord neurons CP1 to CP6 is consistent with earlier studies. **(B)** *tph-1(syb6451)* is expressed in male-specific head neuron class CEM and sex-shared neurons ADF, NSM, and MI. Similar to its expression in hermaphrodites, *tph-1* in MI was previously undetected. In the tail, in addition to previously determined expression in R1B, R3B, and R9B, *tph-1(syb6451)* is also expressed at very low levels in R4B and R7B. Ventral cord expression of *tph-1(syb6451)* in CP1 to CP6 is consistent with previous reports and thus not shown here. **(C)** *bas-1(syb5923)* is expressed in previously identified NSM, ADE, PDE, and CEP neurons. In addition, we detected weak expression in URB as in the hermaphrodite. We also updated *bas-1/AAAD* expression in 39 male-specific neurons and 1 more with variable expression (see **Table S3** for complete list). Neurons are also shown in grayscale for clearer visualization in some cases. Scale bars, 10 μm. Asterisks, non-neuronal expression, also see Fig. 14 and **Fig. S3**.

#### Serotonin/5-HT synthesis

Serotonergic identity had been assigned to several male-specific neurons before (CP1 to CP6, R1B, R3B, R9B)(LOER AND KENYON 1993), and we validated these assignments with our reporter alleles (**Fig. 12**, **Table S3**). In addition, we detected previously unreported expression of *tph-1* (**Fig. 12B**) in the male-specific head neuron class CEM, as well as in a subset of B-type ray sensory neurons, R4B and R7B. However, not all of the neurons display additional, canonical serotonergic neuron features: While R4B and R7B express *bas-1(syb5923)* (with R4B expressing it variably) to generate 5-HT, neither neuron was detected by anti-5-HT staining in the past. On the other hand, R9B and CEM stain positive for 5-HT (Serrano-Saiz *et al*. 2017b), but they do not express *bas-1(syb5923)*, indicating that they may be producing 5-HTP rather than 5-HT (see more below on serotonin uptake). In addition, R4B and R9B, but not R7B or CEM, express *cat-1(syb6486)* for vesicular release of 5-HT.

In the ventral nerve cord, consistent with previous fosmid-based reporter data (Serrano-Saiz *et al*. 2017b), we observed the expression of *cat-1(syb6486)* and *tph-1(syb6451)* in CP1 to CP6 (**Fig. 12A,B**; **Table S3**). Additionally, we also detected novel expression of *bas-1(syb5923)* in CP1 to CP4 and strongly in CP5 and CP6 (**Fig. 12C**, **Table S3**). This updated expression supports the serotonergic identities of these neurons, which had been determined previously based only on their expression of *cat-1/VMAT* reporters and positive staining for 5-HT (LOER AND KENYON 1993; Serrano-Saiz *et al*. 2017b).

#### Dopamine synthesis

We found that the expression of the dopamine-synthesizing *cat-2(syb8255)* reporter allele precisely matched previous assignments of dopaminergic identity (Sulston *et al*. 1975; Sulston *et al*. 1980; LINTS AND EMMONS 1999), i.e. expression was detected exclusively in R5A, R7A, and R9A (**Fig. 13A**, **Table S3**), in addition to all sex-shared dopaminergic neurons. All these neurons show matching expression of *bas-1/AAAD,* the other essential enzyme for dopamine synthesis, and *cat-1/VMAT,* the vesicular transporter for dopamine (**Fig. 13A,C**; **Table S3**).

**Figure 13.**
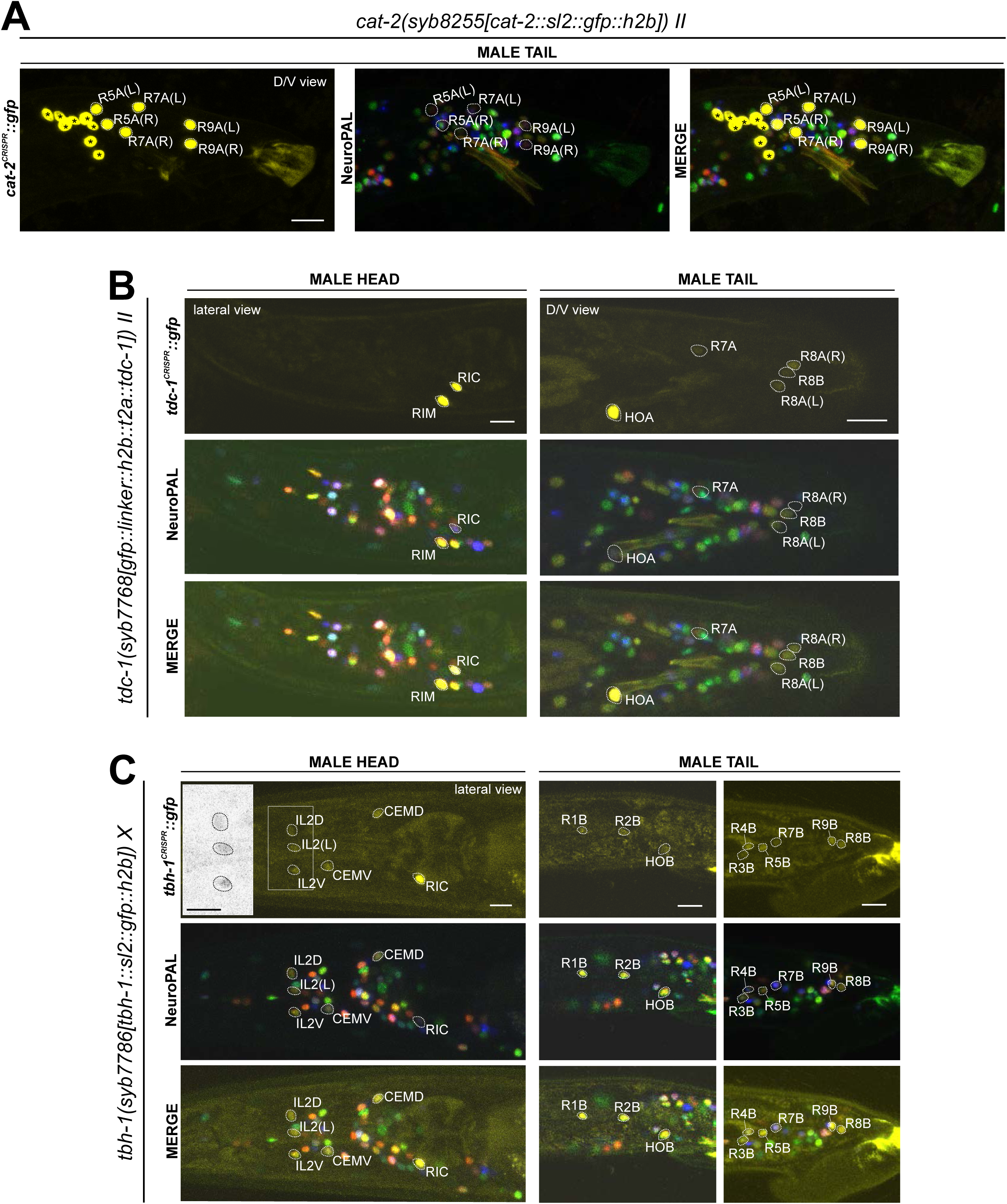
Expression of *cat-2/TH*, *tdc-1/TDC* and *tbh-1/TBH* reporter alleles in the adult male. Neuronal expression was characterized with landmark strain NeuroPAL (*otIs669*). **(A)** *cat-2(syb8255)* is expressed in male-specific neurons R4A, R7A, and R9B. This expression, as well as its expression in sex-shared neurons PDE, CEP, and ADE, is consistent with previous reports (Sulston *et al*. 1975; Sulston *et al*. 1980; LINTS AND EMMONS 1999). **(B)** *tdc-1(syb7768)* is expressed in sex-shared neurons RIM and RIC and male-specific neurons HOA, R8A, and R8B, all consistent with previous studies (Serrano-Saiz *et al*. 2017b). We also detected weak expression in R7A. **(C)** *tbh-1(syb7786)* is expressed in RIC, consistent with its previously reported expression in hermaphrodites. As in hermaphrodites, we also detected *tbh-1(syb7786)* in IL2 neurons of the male. In male-specific neurons, previously unreported expression is detected in CEM, HOB, and all type B ray neurons except for R6B. Intriguingly, this expression pattern resembles that of *pkd-2* and *lov-1*, both genes essential for male mating functions (BARR AND STERNBERG 1999; Barr *et al*. 2001). Inset, grayscale image showing dim expression for IL2 neurons. Scale bars, 10 μm. Asterisks, non-neuronal expression, also see Fig. 14 and **Fig. S3**.

#### Tyramine & Octopamine synthesis

Reporter alleles for the two diagnostic enzymes, *tdc- 1/TDC* and *tbh-1/TBH*, confirm the previously reported assignment of HOA as tyraminergic (Serrano-Saiz *et al*. 2017b), based on the presence of *tdc-1(syb7768)* but absence of *tbh- 1(syb7786)* expression (**Fig. 11B,C**). The *tdc-1* reporter allele reveals a novel site of expression in R7A. Due to lack of *tbh-1* expression, R7A therefore classifies as another tyraminergic neuron. Both HOA and R7A also co-express *cat-1/VMAT* for vesicular release of tyramine.

We detected no neurons in addition to the sex-shared RIC neuron class that shares all features of a functional octopaminergic neuron, i.e. co-expression of *tbh-1/TBH, tdc-1/TDC,* and *cat-1/VMAT.* While one male-specific neuron, R8B, shows an overlap of expression of *tdc- 1(syb7768)* and *tbh-1(syb7786)*, indicating that these neurons can synthesize octopamine, R8B does not express *cat-1(syb6486)*, indicating that these neurons cannot engage in vesicular release of octopamine.

Curiously, while there are no other male-specific neurons that co-express *tdc-1* and *tbh- 1*, several male-specific neurons express *tbh-1*, but not *tdc-1* (**Fig. 13B,C**; **Table 2**, **Table S3**). The absence of the TDC-1/AAAD protein, which produces tyramine, the canonical substrate of the TBH-1 enzyme (**Fig. 1A**), indicates that TBH-1 must be involved in the synthesis of a compound other than octopamine. Moreover, *bas-1/AAAD* is expressed in several of the *tbh- 1*(+); *tdc-1*(-) neurons (R1B, R2B, R3B, R4B, and R7B) (**Fig. 12C**, **Table 2**, **Table S3**). Rather than using L-Dopa or 5-HTP as substrate, BAS-1/AAAD may decarboxylate aromatic amino acids, which then may serve as a substrate for TBH-1. We consider the trace amine phenylethanolamine (PEOH) as a candidate end product (see Discussion).

#### Other monoaminergic neurons

In the preanal ganglion, we detected novel expression of the *cat-1(syb6486)* reporter allele in the cholinergic PDC, PVX, and PVY neurons (**Fig. 12A**). Intriguingly, just as the sex-shared neuron AVL (**Fig. 6C**), these neurons express no other serotonergic, dopaminergic, tyraminergic, or octopaminergic pathway gene. However, we did find PDC (but not PVX or PVY) to express the betaine uptake transporter reporter allele *snf- 3(syb7290)* (**Fig. 9**; more below). PVX and PVY may synthesize or uptake another aminergic transmitter. Such presumptive transmitter is not likely to be synthesized by *hdl-1/AAAD* since we detected no expression of the *hdl-1* reporter allele *syb4208* in the male nervous system (**Fig. S1C,D**).

The expression pattern of the *bas-1/AAAD,* which had not been previously analyzed in the male-specific nervous system, reveals additional novelties. In addition to the “canonical” serotonergic and dopaminergic neurons described above, we detected *bas-1(syb5923)* reporter allele expression in a substantial number of additional neurons, including the tyraminergic HOA and R7A neurons, but also the DVE, DVF, R2A, R3A, R6A, R8A, R2B, R6B, R7B, PCB and SPC neurons (**Fig. 12C**, **Table S3**). As described above, a subset of the neurons co-express *tbh-1(syb7786)* (most B-type ray neurons), a few co-express *tdc- 1(syb7768)* (HOA and several A-type ray neurons), and several co-express neither of these two genes. Only a subset of these neurons express *cat-1(syb6486)*. Taken together, this expression pattern analysis argues for the existence of additional monoaminergic signaling system(s) (**Table 2**).

#### Serotonin/5-HT uptake

In the male-specific nervous system, we detected *mod-5(vlc47)* expression in CEM, PGA, R3B, R9B, and ventral cord neurons CP1 to CP6 (**Fig. 8D**). We found that anti-5-HT staining in CP1 to CP6, R1B, and R3B is unaffected in *mod-5(n3314)* mutant animals, consistent with these neuron expressing the complete 5-HT synthesis machinery (i.e. *tph-1* and *bas-1*)(**Table 2**, **Fig. 8B,D,G**). Hence, like several other monoaminergic neurons, these serotonergic neurons both express, synaptically release, and re-uptake 5-HT. In contrast, anti-5-HT staining is lost from the R9B and PGA neurons of *mod-5(n3314)* mutant animals, indicating that the presence of 5-HT in these neurons depends on 5-HT uptake, consistent with them not expressing the complete 5-HT synthesis pathway (**Table 2**, **Fig. 8B,D,G**). Since R9B and PGA express *cat-1/VMAT,* these neurons have the option to utilize 5-HT for synaptic signaling after *mod-5-*dependent uptake.

#### Tyramine and betaine uptake

We did not observe *oct-1(syb8870)* reporter allele expression in male-specific neurons. As in the hermaphrodite nervous system, we detected *snf-3(syb7290)* in a number of neurons that do not express CAT-1/VMAT (**Table S1**), including in male-specific neurons PHD, and variably, PVV (**Fig. 9C**). As mentioned earlier, the male-specific neuron PDC expresses both *cat-1(syb6486)* and *snf-3(syb7290)*, making it a likely betaine-signaling neuron.

### Sexually dimorphic neurotransmitter expression in sex-shared neurons

#### eat-4/VGLUT

We had previously noted that a fosmid-based *eat-4/VGLUT* reporter is upregulated in the sex-shared neuron PVN, specifically in males (Serrano-Saiz *et al*. 2017b). Since PVN is also cholinergic (**Fig. 4D**)(Pereira *et al*. 2015), this observation indicates a sexually dimorphic co-transmission configuration. As described above (**Fig. 4B**, **Table S2**), our *eat-4* reporter allele revealed low levels of *eat-4/VGLUT* expression in hermaphrodites PVN, but in males the *eat-4* reporter alleles shows strongly increased expression, compared to hermaphrodites. Hence, rather than being an “on” vs. “off” dimorphism, dimorphic *eat-4/VGLUT* expression in male PVN resembles the “scaling” phenomenon we had described previously for *eat-4/VGLUT* in male PHC neurons, compared to hermaphrodite PHC neurons (Serrano-Saiz *et al*. 2017a). Both PHC and PVN display a substantial increase in the amount of synaptic output of these neurons in males compared to hermaphrodites (Cook *et al*. 2019), providing a likely explanation for such scaling of gene expression. The scaling of *eat-4/VGLUT* expression in PVN is not accompanied by scaling of *unc-17/VAChT* expression, which remains comparable in both sexes (**Fig. 4D**).

We also examined AIM, another neuron class that was previously reported to be sexually dimorphic in that AIM expresses *eat-4/VGLUT* fosmid-based reporters in juvenile stages in both sexes, whereas upon sexual maturation its neurotransmitter identity is switched from being glutamatergic to cholinergic only in adult males and not hermaphrodites (Pereira *et al*. 2015; Pereira *et al*. 2019). With the *eat-4(syb4257)* reporter allele, we also detected a downregulation of *eat-4* expression to low levels in young adult males and almost complete elimination in 2-day-old adult males, while expression in hermaphrodites stays high.

#### unc-17/VAChT

The *unc-17/VAChT* reporter allele *syb4491* confirms that cholinergic identity is indeed male-specifically turned on in the AIM neurons (**Fig. 4C**), thereby confirming the previously reported neurotransmitter switch (Pereira *et al*. 2015). The fosmid-based *unc- 17* reporter also showed sexually dimorphic expression in the AVG neurons (Serrano-Saiz *et al*. 2017b). This is also confirmed with the *unc-17* reporter allele, which shows dim and variable expression in hermaphrodites and slightly stronger, albeit still dim, AVG expression in males (**Fig. 4C**, showing a hermaphrodite representing animals with no visible expression and a male with representative dim expression).

#### unc-47/VGAT

*unc-47(syb7566)* confirms previously reported sexually dimorphic expression of *unc-47/VGAT* in several sex-shared neurons, including ADF, PDB, PVN, PHC, AS10, and AS11 (**Fig. 5B**, right side panels) (Serrano-Saiz *et al*. 2017b). The assignment of AS10 was not definitive in our last report (we had considered either DA7 or AS10), but with the help of NeuroPAL the AS10 assignment could be clarified. In all these cases expression was only detected in males and not hermaphrodites. It is worth mentioning that expression of the mCherry-based *unc-47/VGAT* fosmid-based reporter *(otIs564)* in some of these neurons was so dim that it could only be detected through immunostaining against the mCherry fluorophore and not readily visible with the fosmid-based reporter by itself (Serrano-Saiz *et al*. 2017b). In contrast, the *unc-47/VGAT* reporter allele is detected in all cases except the PQR neuron class. In addition, we also detected dim *unc-47/VGAT* expression in the PLM neurons in both sexes (**Fig. 5B**).

#### mod-5/SERT

Expression of the *mod-5(vlc47)* reporter allele is sexually dimorphic in the pheromone-sensing ADF neurons, with higher levels in hermaphrodites compared to males (**Fig. 8F**). Notably, the serotonin-synthesizing enzyme (*tph-1*) and vesicular acetylcholine transporter (*unc-17*) do not exhibit this dimorphism in ADF (**Fig. 8F**). This suggests that the sex difference specifically involves serotonin signaling mechanisms, particularly serotonin uptake rather than synthesis.

We had previously reported that the PVW neuron stains with anti-5-HT antibodies exclusively in males but did not detect expression of a fosmid-based reporter for the serotonin-synthesizing enzyme TPH-1 (Serrano-Saiz *et al*. 2017b). We confirmed the lack of *tph-1* expression with our new *tph-1* reporter allele in both males and hermaphrodites, and also found that hermaphrodite and male PVW does not express the reporter allele for the other enzyme in the 5-HT synthesis pathway, *bas-1.* Because of very dim *cat-1::mCherry* fosmid-based reporter expression that was only detected upon anti-mCherry antibody staining, we had assigned PVW as a 5-HT-releasing neuron (Serrano-Saiz *et al*. 2017b). However, we failed to detect expression of our new *cat-1/VMAT* reporter allele in PVW. Neither did we detect expression of the *mod-5(vlc47)* reporter allele. Taken together, PVW either synthesizes or uptakes 5-HT by unconventional means, akin to the pharyngeal I5 neuron.

In conclusion, although there are some updates in the levels of dimorphic gene expression (PVN and ADF neuron classes), our analysis with reporter alleles does not reveal pervasive novel sexual dimorphism in sex-shared neurons compared to those that we previously identified in (Serrano-Saiz *et al*. 2017b). These sexual dimorphisms are summarized in **Table S4**.

### Neurotransmitter pathway genes in glia

In vertebrates, glia can produce various signaling molecules, including neurotransmitters (Araque *et al*. 2014; SAVTCHOUK AND VOLTERRA 2018). There is some limited evidence for neurotransmitter synthesis in *C. elegans* glia. In males, it had been reported that the socket glia of spicule neurons synthesize and utilize dopamine, based on their expression of *cat-2/TH* and *bas-1/AAAD* (LINTS AND EMMONS 1999; HARE AND LOER 2004; Leboeuf *et al*. 2014). We confirmed this notion with *cat-2/TH* and *bas-1* reporter alleles (**Fig. 14A**). Additionally, we detected expression of *cat-1/VMAT* reporter allele expression in these cells (**Fig. 14A**), indicating that these glia secrete dopamine by canonical vesicular transport. We also observed *bas-1(syb5923)* reporter allele expression in cells that are likely to be the spicule sheath glia (**Fig. 14A**), as well as in additional glia cell types in the head and tail (**Fig. 14B**).

**Figure 14.**
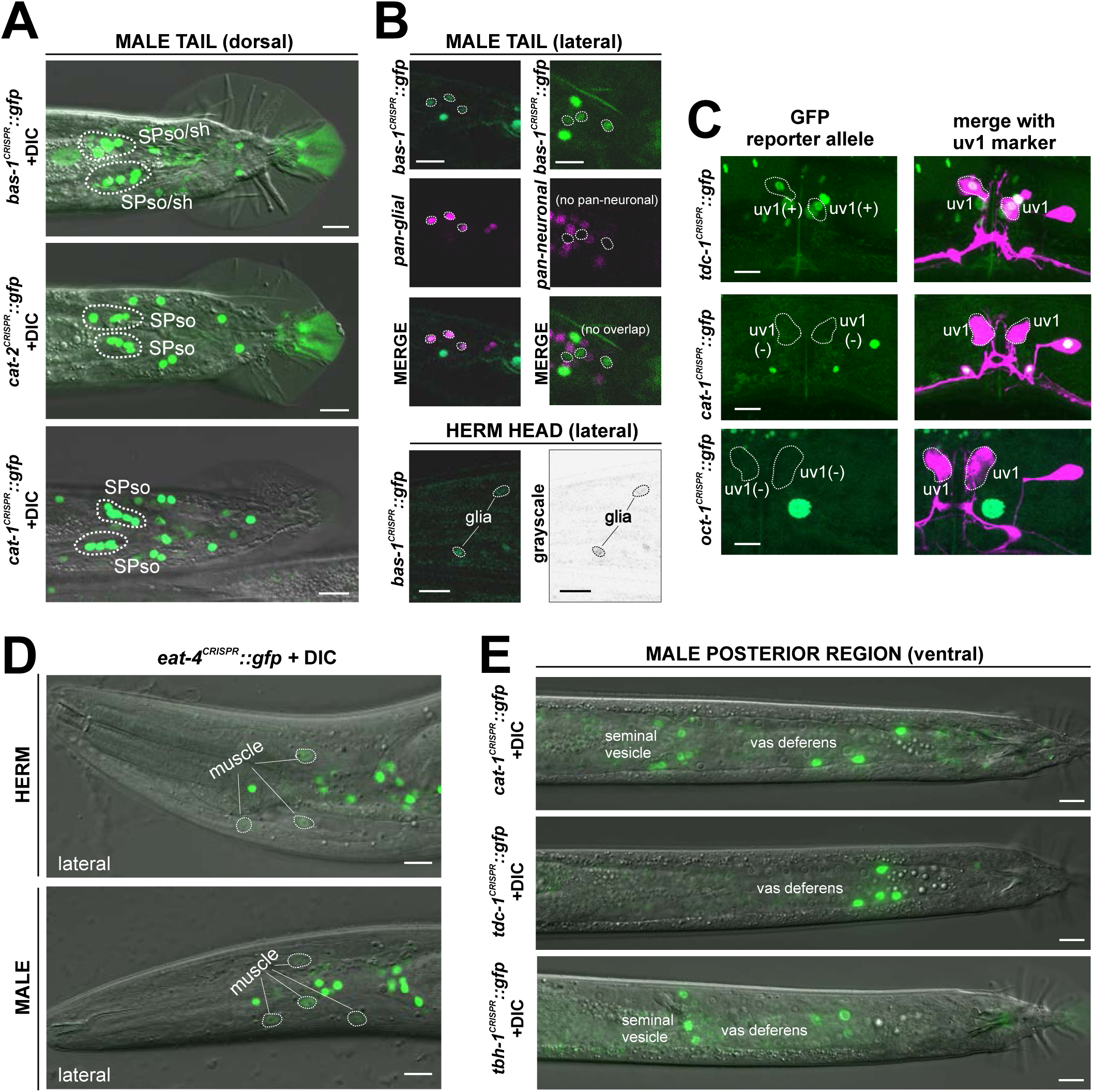
Expression of neurotransmitter pathway genes in non-neuronal cell types. Multiple neurotransmitter pathway genes show expression in glial cells (**A,B**) and other non-neuronal cell types (**C-E**). Also see **Fig. S3** for whole-worm views that capture more non-neuronal expression. **(A)** *bas-1(syb5923)*, *cat-2(syb8255)*, and *cat-1(syb6486)* reporter alleles exhibit expression in the male spicule glial cell types, largely consistent with previous reports.(LINTS AND EMMONS 1999; HARE AND LOER 2004; Leboeuf *et al*. 2014) **(B)** top 6 panels: *bas-1(syb5923)* is expressed in additional, multiple glial cell types in the male tail. Left 3 panels: *bas-1(syb5923)* crossed into a pan-glial reporter *otIs870[mir-228p::3xnls::TagRFP]*, confirming its expression in glial cells; right 3 panels: *bas-1(syb5923)* shows no overlap with the pan-neuronal marker component in NeuroPAL *(otIs669)*. Bottom 2 panels: *bas-1(syb5923)* also shows expression in at least two glial cells in the head. A hermaphrodite head is shown here. Expression is similar in the male. **(C)** In the hermaphrodite vulval region, *tdc-1(syb7768)* is expressed in uv1, consistent with previous reports (Alkema *et al*. 2005). This expression in uv1 is not observed for either *cat-1(syb6486)* or *oct-1(syb8870).* An *ida-1p::mCherry* integrant *vsls269[ida-1::mCherry]* was used for identifying uv1. **(D)** Detection of *eat-4(syb4257)* expression in muscle cells in both sexes, most prominently in the head. **(E)** *cat-1(syb6486)*, *tdc-1(syb7768)*, and *tbh-1(syb7786)* are expressed in the male somatic gonad. All three have expression in the vas deferens; additionally, *cat-1* and *tbh-1* are also expressed in the seminal vesicle.

We detected no expression of other vesicular transporters or neurotransmitter biosynthetic synthesis machinery in glia of either sex. This observation contrasts previous reports on GABA synthesis and release from the AMsh glia cell type (Duan *et al*. 2020; Fernandez-Abascal *et al*. 2022). We were not able to detect signals in AMsh with anti-GABA staining, nor with an SL2 or T2A-based GFP-based reporter allele for any *unc-25* isoform. (Gendrel *et al*. 2016)(M. Gendrel, pers. comm.; this paper).

There is, however, evidence for neurotransmitter uptake by *C. elegans* glial cells, mirroring this specific function of vertebrate glia (HENN AND HAMBERGER 1971). We had previously shown that one specific glia-like cell type in *C. elegans*, the GLR glia, take up GABA via the GABA uptake transporter SNF-11 (Gendrel *et al*. 2016). We did not detect *unc-47/VGAT* fosmid-based reporter expression in the GLRs (Gendrel *et al*. 2016) and also detected no expression with our *unc-47/VGAT* reporter allele. Hence, these glia are unlikely to release GABA via classic vesicular machinery. Other release mechanisms for GABA can of course not be excluded. Aside from the *snf-11* expression in GLR glia (Gendrel *et al*. 2016), we detected expression of the putative tyramine uptake transporter *oct-1/OCT* in a number of head glial cells (**Fig. 8K**), as well as broad glial expression of the betaine uptake transporter *snf-3/BGT1* in the head, midbody, and tail (**Fig. 9E,F**). These results indicate tyramine and betaine clearance roles for glia.

### Neurotransmitter pathway gene expression outside the nervous system

We detected expression of a few neurotransmitter pathway genes in cells outside the nervous system. The most prominent sites of reporter allele expression are located within the somatic gonad. We detected expression of *tdc-1(syb7768)* and *tbh-1(syb7786)* reporter alleles in the gonadal sheath of hermaphrodite as well as *tdc-1(syb7768)* expression in the neuroendocrine uv1 cells (**Fig. 14C; Fig. S3**), as previously reported (Alkema *et al*. 2005). Intriguingly, while *cat-1(syb6486)* is expressed in a midbody gonadal cell posterior to the vulva, likely the distal valve (**Fig. 6C**, **Fig. S3**), we observed no expression of *cat-1(syb6486)* in the gonadal sheath or the uv1 cells (**Fig. 14C**). This suggests alternative release mechanisms for tyramine and octopamine. A vertebrate homolog of the putative tyramine uptake transporter, *oct-1,* has been found to be located presynaptically and to co-purify with synaptosomes (Berry *et al*. 2016; Matsui *et al*. 2016), therefore indicating that this transporter may have the potential to also act in tyramine release, at least in vertebrate cells. However, we observed no expression of our *oct-1* reporter allele in uv1 or gonadal sheath cells.

In the male, *tdc-1(syb7768)*, *tbh-1(syb7786)*, *cat-1(syb6486)*, and *oct-1(syb8870)* animals also show reporter expression in the somatic gonad: while all four genes are expressed in the vas deferens, *cat-1* and *tbh-1*, but not *tdc-1* or *oct-1*, are expressed in the seminal vesicle (**Fig. 14C**, **Fig. 8K**). A similar pattern of *cat-1*(+); *tbh-1*(+); *tdc-1*(-); *oct-1*(-) is detected in several male-specific neurons and may indicate the usage of a novel transmitter (e.g. PEOH, see Discussion) by these cells. *snf-3/BGT1* is also expressed in male somatic gonad cells, indicating that these cells could also use betaine for signaling (**Fig. 9E**).

The AAADs *tdc-1*, as well as *bas-1*, are also prominently expressed in the intestine, where *bas-1* has been shown to be involved in generating 5-HT-derived glucosides (Yu *et al*. 2023). *bas-1*, but not *tdc-1*, is also expressed in the hypodermis and seam cells, as is the betaine uptake transporter *snf-3* (**Fig. S1G, S3**). The *tph-1* reporter allele expresses in a subset of pharyngeal non-neuronal cells during the L1 to L4 larval stages of development (**Fig. S2**), which is consistent with low levels of *tph-1* transcripts detected in pharyngeal muscles in the CeNGEN scRNA dataset. Additionally, we observed previously uncharacterized *eat-4/VGLUT* expression in muscle cells in both sexes (**Fig. 14D**).

## DISCUSSION

Using CRISPR/Cas9-engineered reporter alleles we have refined and extended neurotransmitter assignment throughout all cells of the *C. elegans* male and hermaphrodite. We conclude that in both hermaphrodites and males, about one quarter of neurons are glutamatergic (*eat-4/VGLUT*-positive), a little more than half are cholinergic (*unc-17/VAChT*-positive), around 10% are GABAergic (*unc-25/GAD*-positive), and about another 10% are monoaminergic (*cat-1/VMAT*-positive). We compiled comprehensive lists for gene expression and neuron identities, which are provided in **Table S2** for hermaphrodites and **Table S3** for males. **Figure 3** presents a summary of neurotransmitter usage and atlases showing neuron positions in worm schematics. Additionally, we summarize our rationale for assigning neurotransmitter usage and updates to previously reported data in **Tables 1**, **2**, and **S5**. Given the complexity and nuances in determining neurotransmitter usage, we refer the reader to all the individual tables for a comprehensive description of the subject matter, rather than encouraging sole reliance on the summary in **Figure 3**.

### Neurotransmitter synthesis versus uptake

Direct detection of neurotransmitters through antibody staining has shown that at least two neurotransmitters, GABA and 5-HT, are present in some neurons that do not express the synthesis machinery for these transmitters (**Table 1, 2**). Instead, these neurons acquire GABA and 5-HT through uptaking them via defined uptake transporters, SNF-11/BGT1 for GABA (Mullen *et al*. 2006) and MOD-5/SERT for 5-HT (Ranganathan *et al*. 2001; Jafari *et al*. 2011).

A combination of CeNGEN scRNA transcriptome and our reporter allele data corroborates the absence of synthesis machinery in these presumptive uptake neurons (**Table 1, 2**). One interesting question that relates to these uptake neurons is whether they serve as “sinks” for clearance of a neurotransmitter or whether the taken-up neurotransmitter is subsequently “recycled” for synaptic release via a vesicular transporter. Previous data, as well as our updated expression profiles, provide evidence for both scenarios: ALA and AVF do not synthesize GABA via UNC-25/GAD, but they stain with anti-GABA antibodies in a manner that is dependent on the uptake transporter SNF-11 (Gendrel *et al*. 2016). ALA expresses *unc-47*, hence it is likely to synaptically release GABA, but AVF does not, and it is therefore apparently involved only in GABA clearance. Similarly, RIH, AIM, and PGA express the 5-HT uptake transporter *mod-5/SERT* and stain for 5-HT in a MOD-5-dependent manner (Jafari *et al*. 2011)(this study), but only RIH, not AIM or PGA, expresses the vesicular transporter *cat-1/VMAT*, suggesting RIH is likely a serotonergic signaling neuron whereas AIM and PGA are clearance neurons.

Some neurons do not obviously fall into the synthesis or uptake category, most notably, the anti-GABA-antibody-positive AVA and AVB neurons (both of which conventional cholinergic neurons). None of these neurons express *unc-25/GAD*, nor the *snf-11/BGT1* uptake transporter, yet *unc-25/GAD* is required for their anti-GABA-positive staining (Gendrel *et al*. 2016). This suggests that GABA may be acquired by these neurons through non-canonical uptake or synthesis mechanisms. Also, the AVA and AVB neurons do not express UNC-47 (Gendrel *et al*. 2016; Taylor *et al*. 2021)(this study); hence, it is not clear if or how GABA is released from them. A member of the bestrophin family of ion channels has been shown to mediate GABA release from astrocyte glia in vertebrates (Lee *et al*. 2010) and, more recently, in *C. elegans* (CHENG et al. 2024; GRAZIANO et al. 2024). However, while there are more than 20 bestrophin channels encoded in the *C. elegans* genome (HOBERT 2013), they do not appear to be expressed in the AVA or AVB neurons (Taylor *et al*. 2021).

The co-expression of a specific uptake transporter and a vesicular transporter also leads us to predict the usage of betaine as a potential neurotransmitter. Betaine is known to be synthesized in *C. elegans*, but is also taken up via its diet (Peden *et al*. 2013; Hardege *et al*. 2022). Betaine has documented effects on animal behavior and acts via activation of several betaine-gated ion channels (Peden *et al*. 2013; Hardege *et al*. 2022). Expression of biosynthetic enzymes suggests betaine production in at least the RIM neuron class, which also expresses the vesicular transporter *cat-1/VMAT*, capable of transporting betaine (Hardege *et al*. 2022). The expression of the betaine uptake transporter *snf-3/BGT1* in CAN, AUA, RIR, ASI, and male-specific neuron PDC, coupled with their co-expression of *cat-1/VMAT*, suggests that several distinct neuron classes in different parts of the nervous system may uptake betaine and engage in vesicular betaine release via CAT-1/VMAT to gate betaine-activated ion channels, such as ACR-23 (Peden *et al*. 2013) or LGC-41 (Hardege *et al*. 2022). Additionally, we detected the *snf-3/BGT1* reporter allele in several other neuron classes that do not co-express *cat-1/VMAT*. This indicates that these neurons could function as betaine clearance neurons.

Lastly, based on sequence similarity and expression pattern, we predict that the ortholog of the OCT subclass of SLC22 family, *oct-1*, could serve as a tyramine uptake transporter in *C. elegans*. We identified RIM to be the only neuron expressing an *oct-1* reporter allele, suggesting that like several other monoaminergic neuron classes, RIM both synthesizes its monoaminergic transmitter, tyramine, and reuptakes it after release.

### Evidence for usage of currently unknown neurotransmitters

#### Novel amino acid transmitters?

*unc-47/VGAT* is expressed in a substantial number of non-GABAergic neurons (95 out of 302 total neurons in hermaphrodites, plus 61 out of 93 male-specific neurons). However, expression in many of these non-GABAergic neurons is low and variable and such expression may not lead to sufficient amounts of a functional gene product. Yet, in some neurons (e.g. the SIA neurons) expression of *unc-47* is easily detectable and robust (based on fosmid-based reporter, reporter allele, and scRNA data), indicating that VGAT may transport another presently unknown neurotransmitter (Gendrel *et al*. 2016). In vertebrates, VGAT transports both GABA and glycine, and the same is observed for UNC-47 *in vitro* (Aubrey *et al*. 2007). While the *C. elegans* genome encodes no easily recognizable ortholog of known ionotropic glycine receptors, it does encode anion channels that are closely related by primary sequence (HOBERT 2013). Moreover, a recently identified metabotropic glycine receptor, GPR158 (Laboute *et al*. 2023), has a clear sequence ortholog in *C. elegans, F39B2.8*. Therefore, glycine may also act as a neurotransmitter in *C. elegans.* VGAT has also been shown to transport β-alanine (Juge *et al*. 2013), another potential, but as yet unexplored, neurotransmitter in *C. elegans.* However, it needs to be pointed out that most of the additional *unc-47*-positive neurons do not co-express the LAMP-type UNC-46 protein, which is important for sorting UNC-47/VGAT to synaptic vesicles in conventional GABAergic neurons (Schuske *et al*. 2007). In vertebrates, the functional UNC-46 ortholog LAMP5 is only expressed and required for VGAT transport in a subset of VGAT-positive, GABAergic neurons (Tiveron *et al*. 2016; Koebis *et al*. 2019), indicating that alternative vesicular sorting mechanisms may exist for UNC-47/VGAT.

#### Novel monoaminergic transmitters?

Three neuron classes (AVL, PVX, and PVY) express *cat-1/VMAT* but do not express the canonical synthesis machinery for 5-HT, tyramine, octopamine, or dopamine. Neither do they show evidence for uptake of known monoamines. There are also several *cat-1/VMAT*-positive male-specific neurons that express only a subset of the biosynthetic machinery involved in the biosynthesis of known aminergic transmitters in the worm. That is, some neurons express *cat-1/VMAT* and *bas-1/AAAD*, but none of the previously known enzymes that produce the substrate for BAS-1, i.e. CAT-2 or TPH-1 (**Fig. 1A**). In these neurons, BAS-1/AAAD may decarboxylate an unmodified (i.e. non-hydroxylated) aromatic amino acid as substrate to produce, for example, the trace amine phenylethylamine (PEA) from phenylalanine (**Table 2, Fig. S1A**). A subset of these neurons (all being B-type ray sensory neurons) co-express *tbh-1*, which may use PEA as a substrate to produce the trace amine, phenylethanolamine (PEOH). PEOH is a purported neurotransmitter in Aplysia (Saavedra *et al*. 1977) and the vertebrate brain (SAAVEDRA AND AXELROD 1973) and can indeed be detected in *C. elegans* extracts (F. Schroeder, pers. comm.).

*bas-1/AAAD* may also be responsible for the synthesis of histamine, an aminergic neurotransmitter that can be found in extracts of *C. elegans* (PERTEL AND WILSON 1974). The only other AAAD that displays reasonable sequence similarity to neurotransmitter-producing AAADs is the *hdl-1* gene (HARE AND LOER 2004; HOBERT 2013)(**Fig. S1B**), for which we, however, did not detect any expression in the *C. elegans* nervous system (**Fig. S1D**). Since there are neurons that only express *bas-1/AAAD*, but no enzyme that produces canonical substrates for *bas-1/AAAD* (*tph-1/TPH, cat-2/TH;* **Fig. 1A**) and since at least a subset of these neurons express the monoamine transporter *cat-1/VMAT, bas-1/AAAD* may be involved in synthesizing another currently know bioactive monoamine.

Conversely, based on the expression of *tph-1*, but concurrent absence of *bas-1/AAAD,* the pharyngeal MI neuron, hermaphrodite VC4 and VC5, and male neurons CEM and R9B may produce 5-HTP (**Fig. S1A, Table 2**). 5-HTP may either be used directly as a signaling molecule or it may be metabolized into some other serotonin derivative, an interesting possibility in light of serotonin-derivatives produced elsewhere in the body (Yu *et al*. 2023).

Additionally, 3 neuron classes (IL2, HOB, and R5B) express *tbh-1* but lack expression of any other genes in canonical monoaminergic pathways, including *bas-1* (**Table 2**). This observation further suggests the presence of non-canonical mechanisms for monoaminergic synthesis. Taken together, monoaminergic pathway genes are expressed in unconventional combinations in several neuron classes, pointing towards the existence of yet undiscovered monoaminergic signaling systems.

### Neurons devoid of canonical neurotransmitter pathway genes may define neuropeptide-only neurons

We identified neurons that do not express any conventional, well-characterized vesicular neurotransmitter transporter families, namely UNC-17/VAChT, CAT-1/VMAT (the only SLC18 family members), UNC-47/VGAT (only SLC32 family member), or EAT-4/VGLUT (an SLC17 family member). Six sex-shared neurons (AVH, BDU, PVM, PVQ, PVW, RMG) and one male-specific neuron (SPD) fall into this category. Most of these neurons exhibit features that are consistent with them being entirely neuropeptidergic. First, electron microscopy has revealed a relative paucity of clear synaptic vesicles in most of these neurons (White *et al*. 1986; Cook *et al*. 2019; Witvliet *et al*. 2021). Second, not only do these neurons express a multitude of neuropeptide-encoding genes (Taylor *et al*. 2021), but they also display a dense interconnectivity in the “wireless” neuropeptidergic connectome (Ripoll-Sanchez *et al*. 2022).

That said, electron microscopy shows that some of the neurons devoid of conventional neurotransmitter pathway genes generate synapses with small, clear synaptic vesicles, indicative of the use of non-peptidergic transmitters (e.g. the sex-shared RMG and PVM neurons or the male-specific SPD neurons) (White *et al*. 1986; Cook *et al*. 2019; Witvliet *et al*. 2021). It is therefore conceivable that either conventional neurotransmitters utilize non-conventional neurotransmitter synthesis and/or release pathways, or that completely novel neurotransmitter systems remain to be discovered. Although the *C. elegans* genome does not encode additional members of the SLC18A2/3 (*cat-1/VMAT, unc-17/VAChT*) or SLC32A1 (*unc-47/VGAT*) family of vesicular neurotransmitter transporters, it does contain a number of additional members of the SLC17A6/7/8 (VLGUT) family (HOBERT 2013). These may serve as non-canonical vesicular transporters of more uncommon neurotransmitters or, alternatively, may be involved in modulating release of glutamate (Serrano-Saiz *et al*. 2020; Choi *et al*. 2021). Uncharacterized paralogs of *bona fide* neurotransmitter uptake transporters (SLC6 superfamily) may also have functions in neurotransmitter release rather than uptake. However, based on CeNGEN scRNA data, no robust or selective expression of these SLC17 or SLC6 family members is observed in these “orphan neurons”.

### Co-transmission of multiple neurotransmitters

Our analysis expands the repertoire of neurons that co-transmit multiple neurotransmitters (**Fig. 3**). Neurotransmitter co-transmission has been observed in multiple combinations in the vertebrate brain (WALLACE AND SABATINI 2023). In *C. elegans,* the most frequent co-transmission configurations are a classic, fast transmitter (acetylcholine or glutamate) with a monoamine. Co-transmission of two distinct monoaminergic systems also exists. In several cases, however, it is not clear whether the second neurotransmitter is indeed used for communication or whether its presence is merely a reflection of this neuron being solely an uptake neuron. For example, the glutamatergic AIM neuron stains positive for serotonin, which it uptakes via the uptake transporter MOD-5, but it does not express the vesicular monoamine transporter *cat-1/VMAT* (**Fig. 3, 6, 8**, **Table 1**, **2**).

Co-transmission of small, fast-acting neurotransmitters (glutamate, GABA, acetylcholine) does exist, but it is rare (**Fig. 3**). The most prominent co-transmission configuration is acetylcholine with glutamate, but acetylcholine can also be co-transmitted with GABA. There are no examples of co-transmission of glutamate and GABA, as observed in several regions of the vertebrate brain (WALLACE AND SABATINI 2023).

Interestingly, co-transmission appears to be much more prevalent in the male-specific nervous system, compared to the sex-shared nervous system (**Fig. 3**, **Table S3**). Remarkably, several male-specific neuron classes may utilize three co-transmitters. Such extensive co-transmission may relate to male-specific neurons displaying a greater degree of anatomical complexity compared to the hermaphrodite nervous system, both in terms of branching patterns and extent of synaptic connectivity (Jarrell *et al*. 2012; Cook *et al*. 2019). Given that all co-transmitting neurons display multiple synaptic outputs (Cook *et al*. 2019), it appears possible that each individual neurotransmitter secretory system is distributed to distinct synapses. Based on vertebrate precedent (WALLACE AND SABATINI 2023), co-release from the same vesicle is also possible.

### Sexual dimorphisms in neurotransmitter usage

The observation of sexual dimorphisms in neurotransmitter abundance in specific regions of the mammalian brain has been one of the earliest molecular descriptors of neuronal sex differences in mammals (Mccarthy *et al*. 1997). However, it has remained unclear whether such differences are the result of the presence of sex-specific neurons or are indications of distinctive neurotransmitter usage in sex-shared neurons. Using *C. elegans* as a model, we have been able to precisely investigate (a) whether sex-specific neurons display a bias in neurotransmitter usage and (b) whether there are neurotransmitter dimorphisms in sex-shared neurons (Pereira *et al*. 2015; Gendrel *et al*. 2016; Serrano-Saiz *et al*. 2017b)(this paper). We found that male-specific neurons display a roughly similar proportional usage of individual neurotransmitter systems and note that male specific neurons display substantially more evidence of co-transmission, a possible reflection of their more elaborate morphology and connectivity. We also confirmed evidence for sexual dimorphisms in neurotransmitter usage in sex-shared neurons (**Table S4**), which are usually correlated with sexual dimorphisms in synaptic connectivity of these sex-shared neurons (Cook *et al*. 2019).

### Neurotransmitter pathway genes in glia and gonad

Neurotransmitter uptake is a classic function of glial cells across animal phylogeny (HENN AND HAMBERGER 1971), and such uptake mechanisms are observed in *C. elegans* as well. Previous reports demonstrated glutamate uptake by CEPsh (Katz *et al*. 2019) and GABA uptake by GLR glia (Gendrel *et al*. 2016). We now add to this list betaine uptake by most glia, as inferred from the expression pattern of SNF-3/BGT1 (**Fig. 9**, **Table S1**).

Studies in vertebrates have also suggested that specific glial cell types synthesize and release several neurotransmitters (Araque *et al*. 2014; SAVTCHOUK AND VOLTERRA 2018). For example, astrocytes were recently shown to express VGLUT1 to release glutamate (DE Ceglia *et al*. 2023). Evidence of neurotransmitter synthesis and release also exists in *C. elegans.* Previous work indicated that glia associated with male-specific spicule neurons synthesize (through *cat-2/TH* and *bas-1/AAAD*) the monoaminergic transmitter dopamine to control sperm ejaculation (Leboeuf *et al*. 2014). Our identification of *cat-1/VMAT* expression in these glia indicate that dopamine is released via the canonical vesicular monoamine transporter. We also detected expression of *bas-1/AAAD* in additional male and hermaphrodite glia, indicating the production of other signaling substances released by these glia. *bas-1* has indeed recently been shown to be involved in the synthesis of a class of unconventional serotonin derivates (Yu *et al*. 2023).

We observed no additional examples of neurotransmitter synthesis and release by glia, based on the apparent absence of detectable expression of neurotransmitter-synthesizing enzymes or any vesicular transporter (*unc-17/VAChT, unc-47/VGAT, eat-4/VGLUT, cat-1/VMAT*). Both observations are particularly notable in the context of previous reports on GABA synthesis and release from the AMsh glia cell type (Duan *et al*. 2020; Fernandez-Abascal *et al*. 2022). We were not able to detect AMsh with anti-GABA staining, nor with reporter alleles of *unc-25/GAD*. However, since very low levels of *unc-25* are observed in AMsh scRNA datasets (Taylor *et al*. 2021; Purice *et al*. 2023), the abundance of GABA in AMsh may lie below conventional detection levels.

Outside the nervous system, the most prominent and functionally best characterized usage of neurotransmitters lies in the hermaphrodite somatic gonad, which has been shown to synthesize octopamine and use it to control oocyte quiescence (Alkema *et al*. 2005; Kim *et al*. 2021). Intriguingly, we also detected *tbh-1, tdc-1,* and *cat-1* expression in the somatic gonad of the male, specifically the vas deferens, which is known to contain secretory granules that are positive for secretory molecular markers (Nonet *et al*. 1993). The presence of octopamine is unexpected because, unlike oocytes, sperm are not presently known to require monoaminergic signals for any aspect of their maturation. It will be interesting to assess sperm differentiation and function of *tbh-1* or *tdc-1* mutant animals. The usage of monoaminergic signaling systems in the gonad is not restricted to *C. elegans* and has been discussed in the context of sperm functionality and oocyte maturation in vertebrates (Mayerhofer *et al*. 1999; Ramirez-Reveco *et al*. 2017; Alhajeri *et al*. 2022).

### Comparing approaches and caveats of expression pattern analysis

Our analysis also provides an unprecedented and systematic comparison of antibody staining, CeNGEN scRNA transcript data, reporter transgene expression, and knock-in reporter allele expression. The bottom-line conclusions of these comparisons are: (1) Reporter alleles reveal more sites of expression than fosmid-based reporters. It is unclear whether this is due to the lack of *cis*-regulatory elements in fosmid-based reporters or issues associated with the multicopy-nature of these reporters (e.g. RNAi-based gene silencing of multicopy arrays or squelching of regulatory factors). Another factor to consider is that neuron identification for most fosmid-based reporters was carried out prior to the introduction of NeuroPAL. Consequently, errors occasionally occurred, as exemplified by the misidentification of neuron IDs for CA7 and CP7 in previous instances (Serrano-Saiz *et al*. 2017b). (2) The best possible reporter approaches (i.e. reporter alleles) show very good overlap with scRNA data, thereby validating each approach. However, our comparisons also show that no single approach is perfect. CeNGEN scRNA data can miss transcripts and can also show transcripts in cells in which there is no independent evidence for gene or protein expression. Conversely, antibody staining displays vagaries related to staining protocols and protein localization, which can be overcome with reporter approaches, but the price to pay with reporter alleles is that if they are based on SL2 or T2A strategies, they may fail to detect additional levels of posttranslational regulation, which may result in protein absence even in the presence of transcripts. The existence of such mechanisms may be a possible explanation for cases where the expression of synthesis and/or transport machinery expression does not match up (e.g. *tdc-1*(-); *tbh-1*(+) neurons).

Our detailed analysis of reporter allele expression has uncovered several cases where expression of a neurotransmitter pathway gene in a given neuron class appears very low and variable from animal to animal. Such variability only exists when expression is dim, thus one possible explanation for it is that expression levels merely hover around an arbitrary microscopical detection limit. However, we cannot rule out the other possibility that this may also reflect true on/off variability of gene expression. Taking this notion a step further, we cannot exclude the possibility that expression observed with reporter alleles misses sites of expression. This possibility is raised by our inability to detect *unc-25/GAD* reporter allele expression in AMsh glia (Duan *et al*. 2020; Fernandez-Abascal *et al*. 2022) or *eat-4* reporter allele expression in AVL and DVB neurons, in which some (but not other) multicopy reporter transgenes revealed expression of the respective genes (Li *et al*. 2023). Functions of these genes in the respective cell types were corroborated by cell-type specific RNAi experiments and/or rescue experiments; whether there is indeed very low expression of these genes in those respective cells or whether drivers used in these studies for knock-down and/or rescue produce very low expression in other functionally relevant cells remains to be resolved.

## Conclusions

In conclusion, we have presented here the most complete neurotransmitter map that currently exists for any animal nervous system. Efforts to map neurotransmitter usage on a system-wide level are well underway in other organisms, most notably, *Drosophila melanogaster* (DENG et al. 2019; ECKSTEIN et al. 2024). The *C. elegans* neurotransmitter map presented here comprises a critical step toward deciphering information flow in the nervous system and provides valuable tools for studying the genetic mechanisms underlying cell identity specification. Moreover, this neurotransmitter map opens new opportunities for investigating sex-specific neuronal differentiation processes, particularly in the male-specific nervous system, where a scarcity of molecular markers has limited the analysis of neuronal identity control. Lastly, our analysis strongly suggests that additional neurotransmitter systems remain to be identified.

While the gene expression patterns delineated here enable informed predictions about novel neuronal functions and neurotransmitter identities, further investigations involving genetic perturbations, high-resolution imaging, complementary functional assays, and analyses across developmental stages are needed to shed further light on neurotransmitter usage. Nonetheless, this comprehensive neurotransmitter map provides a robust foundation for deciphering neural information flow, elucidating developmental mechanisms governing neuronal specification, exploring sexual dimorphisms in neuronal differentiation, and potentially uncovering novel neurotransmitter systems awaiting characterization.

## RESOURCE AVAILABILITY

### Lead contact

Further information and requests for resources and reagents should be directed to and will be fulfilled by the Lead Contact, Oliver Hobert.

### Materials availability

All newly generated strains are available at the Caenorhabditis Genetics Center (CGC).

### Data and code availability

Any additional information required to analyze the data reported in this paper is available from the lead contact upon request.

## ACKNOWLEDGEMENTS

We thank Chi Chen for generating nematode strains. We thank Emily Bayer, James Rand, Piali Sengupta, and Esther Serrano-Saiz for comments on the manuscript, Frank Schroeder and Marie Gendrel for discussion and communicating unpublished results, Aakanksha Singhvi for discussing glia scRNA data and Michael Koelle for an *ida-1* reporter strain. Some strains were provided by the CGC, which is funded by NIH Office of Research Infrastructure Programs (P40 OD010440).

## FUNDING

This work was funded by the Howard Hughes Medical Institute and by NIH R01 NS039996.

## CONFLICT OF INTEREST

The authors declare no conflicts of interest.

## SUPPLEMENTARY FIGURE AND TABLE LEGENDS

**Figure S1. Use of AAADs (Aromatic Amino Acid Decarboxylases) in *C. elegans*.**

(**A**) Biosynthesis of biogenic amines involve the use of AAADs. Modified from (HOBERT 2013)

(**B**) Phylogenetic trees of amino acid decarboxylases. The only AAAD that displays reasonable sequence similarity to neurotransmitter-producing AAADs is the *hdl-1* gene (HARE AND LOER 2004; HOBERT 2013).

(**C**, **D**) We engineered a GFP reporter allele for *hdl-1* (*syb1048*) (**C**) and did not detect any expression (**D**). We also attempted but failed at amplifying weak expression signals by using a Cre recombination strategy (**C**, *syb4208,* see Methods).

**Figure S2. *tph-1/TPH* reporter allele expression in the hermaphrodite larvae.** Hermaphrodite heads from different larval stages (L1 to L4) and young adults expressing *tph-1(syb6451). tph-1* expression in the NSML/R and ADFL/R neuron pairs and in the MI neuron was visible across all larval stages and during adulthood. MI expression was validated using *otIs518[eat-4(fosmid)::SL2::mCherry::H2B]*, a reporter for the glutamatergic identity of MI. Non-neuronal expression of *tph-1* (asterisks) could be detected in a subset of pharyngeal muscles in the L1 to L4 larval stages but very dim or no expression was detected in young adults. Scale bars, 10 μm.

**Figure S3. Whole-worm images showing monoaminergic pathway gene expression in different tissue types.** Monoaminergic neurotransmitter reporters show abundant expression outside of the nervous system. Lateral views of entire worms expressing the *tph-1/TPH* (*syb6451*), *bas-1/AAAD* (*syb5923*), *cat-2/TH* (*syb8255*), *cat-1/VMAT* (*syb6486*), *tdc-1/TDC* (*syb7768*), and *tbh-1/TBH* (*syb7786*) reporter alleles.

(**A**) GFP and DIC views.

(**B**) Grayscale views of the GFP signal with tissue types labeled as noted on the figure. Scale bars, 20 μm. For more details, see Fig. 14.

**Table S1. scRNA data for neurotransmitters in the hermaphrodite.** Here we show expression of previous reporters and reporter alleles used in this study, compared to scRNA data. Note that scRNA expression values for *eat-4* and *unc-47* can be unreliable because they were overexpressed to isolate individual neurons for scRNA analysis (Taylor *et al*. 2021).

**Table S2. Updated expression patterns of neurotransmitter pathway genes in hermaphrodites.**

**Table S3. Updated expression patterns of neurotransmitter pathway genes in male-specific neurons.**

**Table S4. Summary of sexually dimorphic use of neurotransmitter pathway genes in sex-shared neurons.**

**Table S5. Summary of updates to expression patterns of classic neurotransmitter pathway genes.**

## Notes

### Competing Interest Statement

The authors have declared no competing interest.

### Summary of Updates

Response to reviewers comments have been incorporated. Several new reporter alleles for other neurotransmitter pathway genes have been added.

