## Supplementary figures and images for "A neurotransmitter atlas of *C. elegans* males and hermaphrodites"

### Fig2#

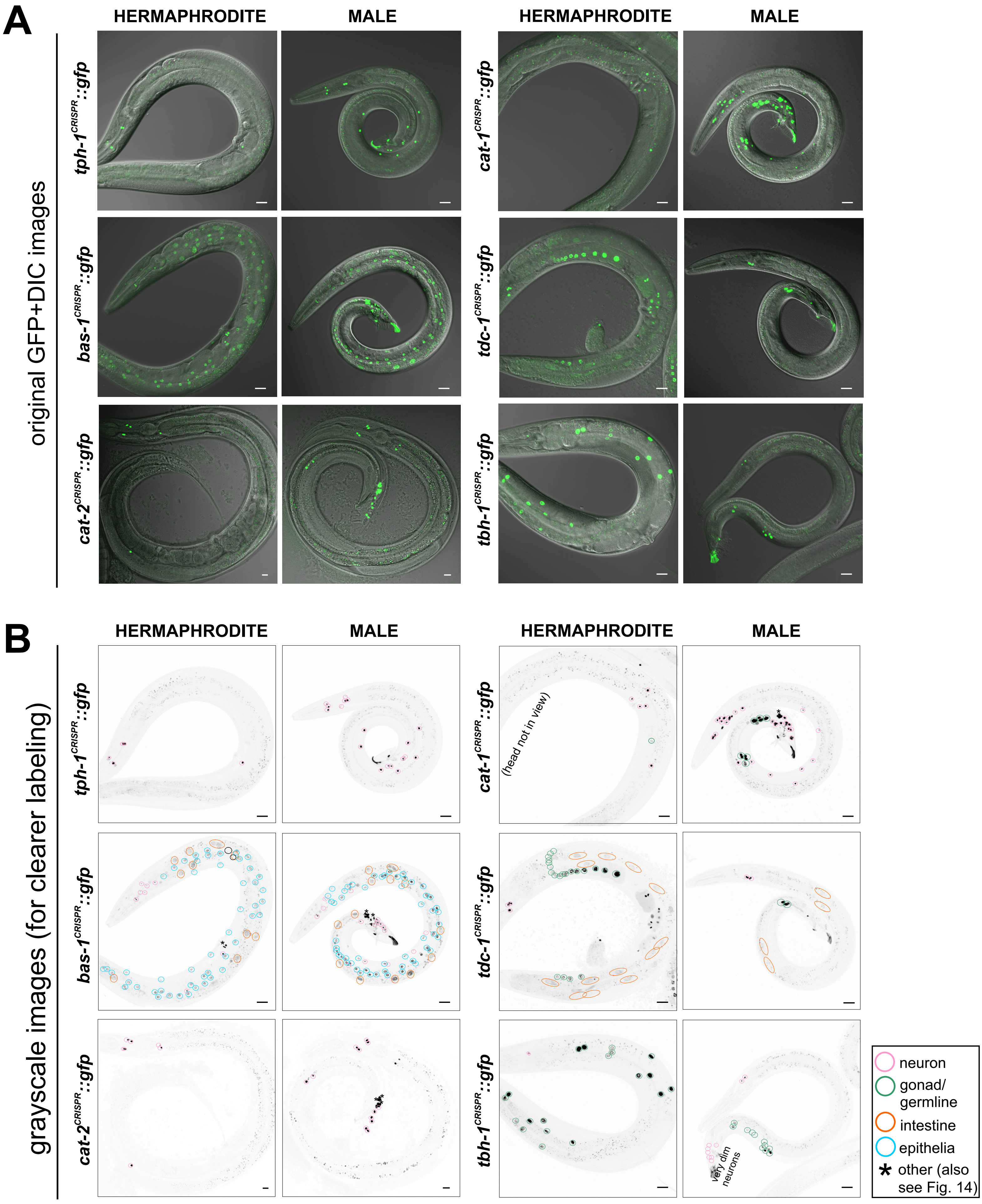

### FigS1

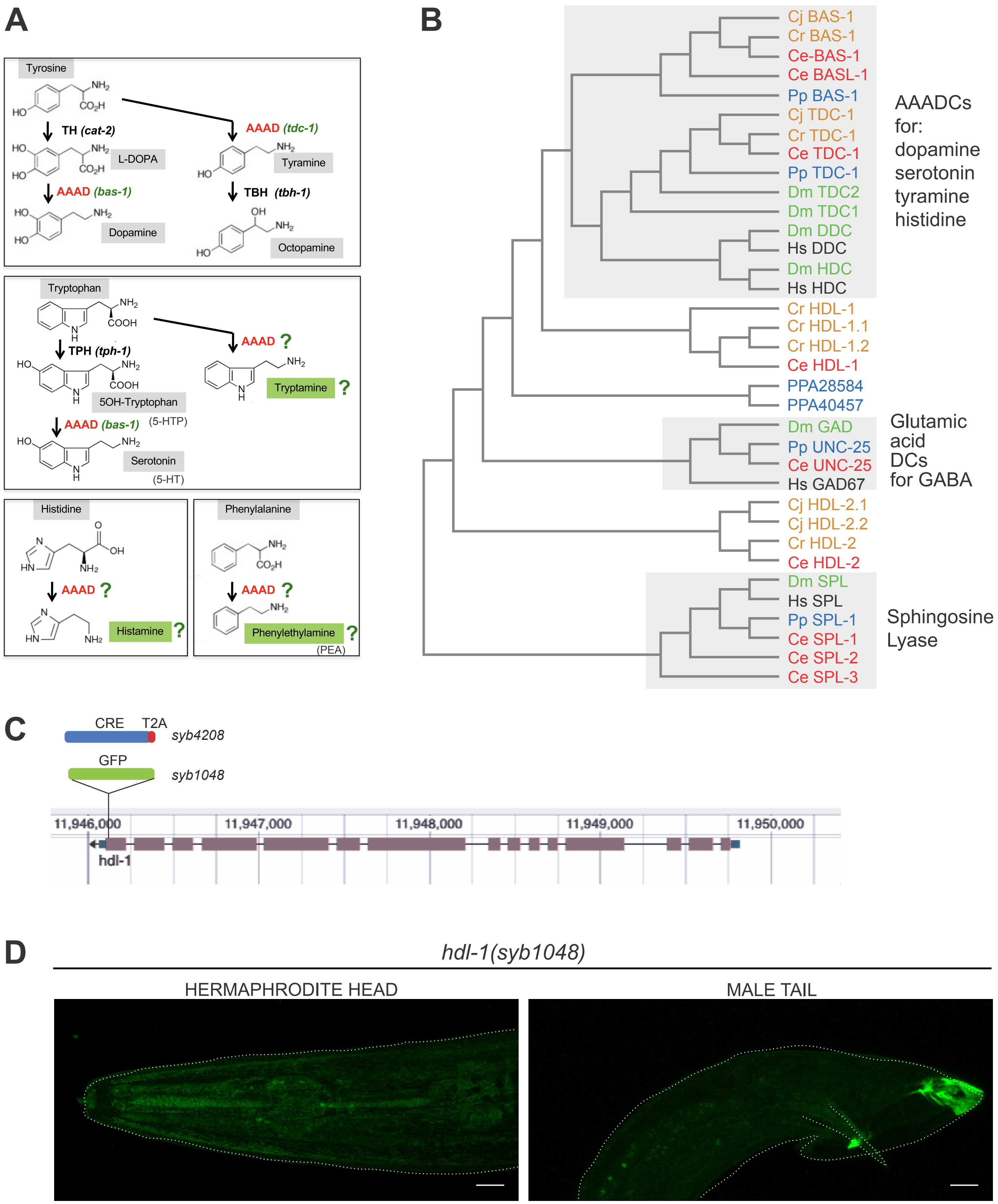

### FigS2

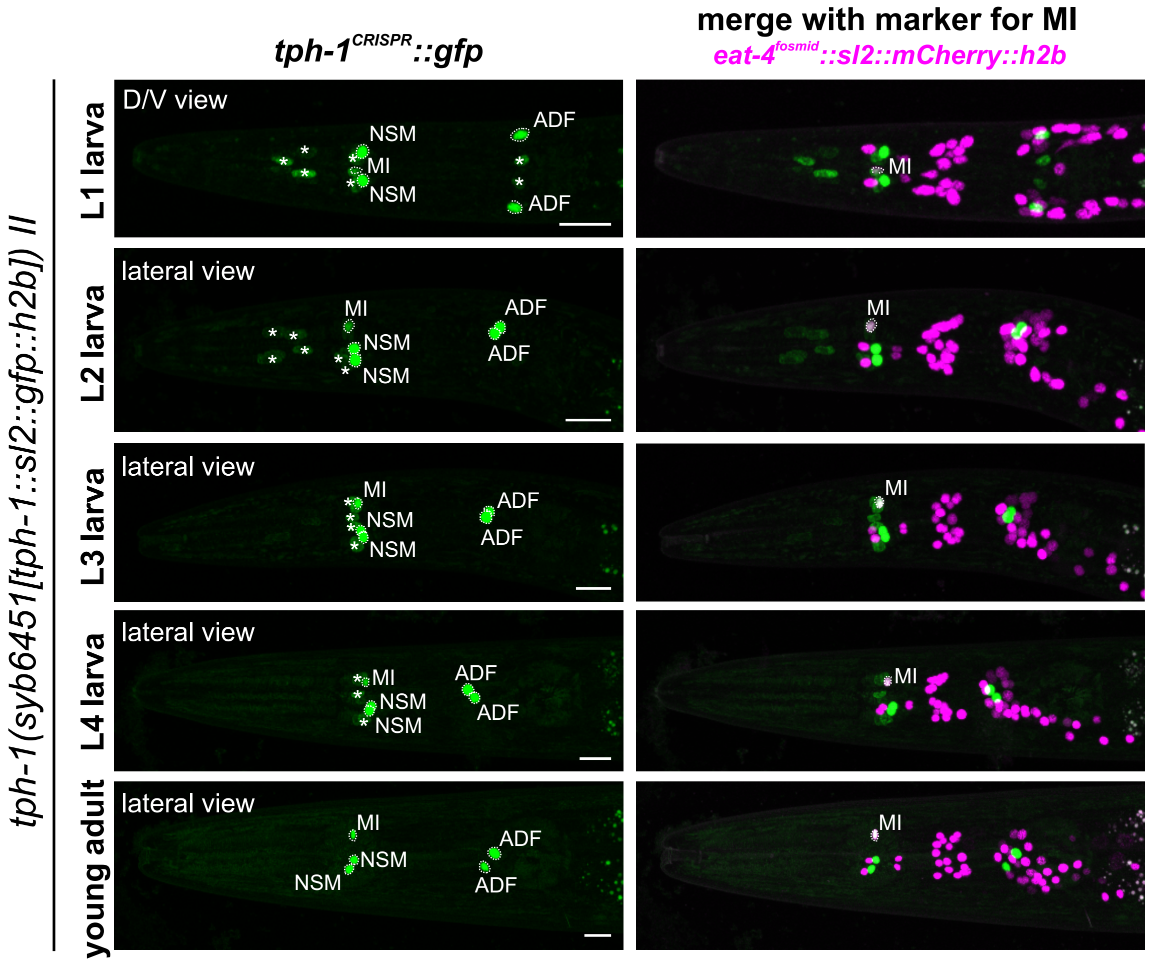
